# Impact of deprivation and preferential usage on functional connectivity between early visual cortex and category selective visual regions

**DOI:** 10.1101/2024.05.17.593020

**Authors:** Leland L. Fleming, Matthew Defenderfer, Pinar Demirayak, Paul Stewart, Dawn K. Decarlo, Kristina M. Visscher

## Abstract

Human behavior can be remarkably shaped by experience, such as the removal of sensory input. Many studies of conditions such as stroke, limb amputation, and vision loss have examined how the removal of input changes brain function. However, an important question has yet to be answered: when input is lost, does the brain change its connectivity to preferentially use some remaining inputs over others? In individuals with healthy vision, the central portion of the retina is preferentially used for everyday visual tasks, due to its ability to discriminate fine details. However, when central vision is lost in conditions like macular degeneration, peripheral vision must be relied upon for those everyday tasks, with certain portions receiving “preferential” usage over others. Using resting-state fMRI collected during total darkness, we examined how deprivation and preferential usage influence the intrinsic functional connectivity of sensory cortex by studying individuals with selective vision loss due to late stages of macular degeneration. We found that cortical regions representing spared portions of the peripheral retina, regardless of whether they are preferentially used, exhibit plasticity of intrinsic functional connectivity in macular degeneration. Cortical representations of spared peripheral retinal locations showed stronger connectivity to MT, a region involved in processing motion. These results suggest that long-term loss of central vision can produce widespread effects throughout spared representations in early visual cortex, regardless of whether those representations are preferentially used. These findings support the idea that connections to visual cortex maintain the capacity for change well after critical periods of visual development.

**Highlights:**

- Portions of early visual cortex representing central vs. peripheral vision exhibit different patterns of connectivity to category-selective visual regions.
- When central vision is lost, cortical representations of peripheral vision display stronger functional connections to MT than central representations.
- When central vision is lost, connectivity to regions selective for tasks that involve central vision (FFA and PHA) are not significantly altered.
- These effects do not depend on which locations of peripheral vision are used more.

## Introduction

In 1890, the great American psychologist William James noted that “Organic matter, especially nervous tissue, seems endowed with a very extraordinary degree of plasticity” (James, 1890). Since then, the field of neuroscience has sought to understand the ways in which the brain is shaped by experience. This question has been of particular interest in understanding the nature of sensory systems, whose development and function rely heavily on experience (Chang and Merzenich, 2003; Grubb and Thompson, 2004; Hubel and Wiesel, 1962; Hubel and Wiesel, 1963; Zhang et al., 2001). Seminal work by Hubel and Wiesel demonstrated the effects of experience on the developing visual system (Hubel and Wiesel, 1962), revealing that monocular deprivation early in life blocks the formation of ocular dominance columns. Other studies have examined the relationship between sensory deprivation and plasticity by observing cortical changes following retinal lesions in adult animals (Calford et al., 2000; Gilbert and Li, 2012; Gilbert and Wiesel, 1992; Kaas et al., 1990). At a larger scale, deprivation of input has been shown to influence connection patterns between retinotopically mapped cortical regions and other parts of cortex (Plank et al., 2017; Sabbah et al., 2017). It is important to note, however, that experiences like sensory deprivation are often multifaceted and can involve multiple, simultaneous features (Cramer et al., 2011; Liu et al., 2010; Toyoizumi et al., 2014; Turrigiano, 1999). For example, the experience of partial sensory deprivation can include deprivation of input to some brain regions, maintenance of input to spared regions, and preferential, increased usage of regions of cortex associated with spared sensory input (Liu et al., 2010). Here we separate the impact of these different features of experience, in a human model examining connections between early visual cortical areas and category-selective cortical areas.

Studying patients with selective sensory impairment, such as central vision loss due to late-stage macular degeneration (MD), can inform our understanding of how the brain adapts to changes in experience (Baker et al., 2005; Baker et al., 2008; Baseler et al., 2011; Burge et al., 2016; Dilks et al., 2009; Masuda et al., 2008; Masuda et al., 2021; Sabbah et al., 2017; Sunness et al., 2004). In later stages of MD, photoreceptor cells degenerate, forming a lesion in the center of the retina (Zarbin, 2004). This results in the deprivation of bottom-up, retinal input to the cortical representation of the lesioned area -- the so-called lesion projection zone (LPZ). Due to the lack of central vision, many individuals with MD preferentially use a specific part of their peripheral vision, known as the “preferred retinal locus” (PRL), as an oculomotor reference point for everyday visual tasks like recognizing faces (Bullimore et al., 1991) and reading (Timberlake et al., 1987). The preferential usage of the PRL makes it functionally distinct from other “un-preferred retinal loci” (URLs), which maintain sensory input to visual cortex, but do not necessarily receive increased use. Due to the precise retinotopy of the visual cortex, projections from PRL and URL regions of the retina can be predicted based on anatomy (Benson et al., 2012) and the location of those projections is not thought to change substantially after MD (Baseler et al., 2011). However, little is known about how the cortical representations of the PRL and URL (referred to here as the cPRL and cURL) are differentially altered in response to input deprivation or preferential usage.

Functional connectivity, or the correlation in spontaneous activity between brain regions (Biswal et al., 1995), is a stable and reproducible metric (Gratton et al., 2018), used to assess changing brain function in many contexts, including development (Dosenbach et al., 2010; Fair et al., 2007; Satterthwaite et al., 2013), in aging (Andrews-Hanna et al., 2007; Chan et al., 2014; Geerligs et al., 2015; Meunier et al., 2009), as well as with changes in visual experience (Lewis et al., 2009). Because central and peripheral vision are used differently after central vision loss, this loss offers an opportunity to examine alterations in typical patterns of functional connectivity in visual cortex. Measuring spontaneous fluctuations in brain activity at rest provides a window into the intrinsic properties of brain function (Biswal et al., 1995; Fox and Raichle, 2007; Raichle et al., 2001). Changes in functional connectivity brought on by experience may reflect a history of repeated co-activation of brain regions over time in a Hebbian-like fashion (Harmelech and Malach, 2013; Hebb, 1949). Previous work shows that functional connectivity in primary visual cortex is retinotopically organized so that regions that represent central vision have different functional connectivity patterns from those that represent peripheral vision (Arcaro et al., 2015; Genç et al., 2016; Griffis et al., 2017; Heinzle et al., 2011; Raemaekers et al., 2014; Striem-Amit et al., 2015). These differences are thought to reflect the differential functions of central and peripheral vision. Thus, examining patterns of functional connections in people with central vision loss due to macular degeneration, where peripheral vision must be used for all visual tasks, provides a window to understand plasticity of a key organizing feature of visual cortex.

Prior work suggests that functional connections change following vision loss. For example, following the loss of central vision, cortical representations of peripheral vision in early visual cortex exhibit altered patterns of functional connectivity to ventral occipital cortex (Sabbah et al., 2017). This finding is noteworthy given that ventral occipital cortex exhibits different patterns of connectivity to central versus peripheral representations in early visual cortex (Park et al., 2018). These differential patterns of connectivity for central versus peripheral representations likely reflect the ways in which central and peripheral vision are normally used for different types of visual tasks. For example, central vision is typically used more for everyday visual tasks such as recognizing faces. Thus, cortical representations of central vision are more strongly associated with regions with high selectivity for faces, like fusiform face area (Levy et al., 2001). On the other hand, there may be specific roles for peripheral vision that are associated with other category selective regions. For example, human area MT is involved in identifying looming or moving stimuli (Tootell et al., 1995). While motion detection is important throughout the visual field, it is performed particularly well with peripheral vision (Yu et al., 2010). Here, we directly test the prediction that changes in functional connectivity associated with increased usage of peripheral vision are specific to the region that is preferentially used (i.e. the preferred retinal locus).

To investigate this hypothesis, we examined how patterns of spontaneous activity in the complete absence of visual input are changed in MD. We focused on functional connectivity between early visual cortex and higher-order visual regions that exhibit selectivity for specific types of visual stimuli. In doing so, we seek to understand how different features of experience, in the form of deprivation, maintenance, and preferential usage produce macroscale changes in the intrinsic properties of visual cortex that are likely outcomes of microscale forms of plasticity.

## Materials and Methods

### Participants

Patients were recruited as part of the Connectomes in Human Diseases MD Plasticity project. Inclusion criteria for MD patients for the project were: 1) central vision loss in both eyes for a minimum of two years, 2) a clearly-defined preferred retinal locus as determined from a Macular Integrity Assessment (MAIA) microperimetry, 3) visual acuity of 20/100 or worse in their best eye, and 4) have a matched healthy control participant. Retinal microperimetry and a visual acuity assessment were performed in order to verify that each MD participant had significant loss of central vision. Visual acuity was measured using the Early Treatment Diabetic Retinopathy Study (ETDRS) test (Ferris et al., 1982).

Healthy control participants were matched on the basis of gender, age (matched to an individual participant +/- 5 years), and education level (no high school diploma, high school diploma, some college, college degree, or advanced degree). Control participants were required to have normal or corrected-to-normal vision and be free of ocular disease.

The data analyzed here includes 11 MD patients (5 males, 6 females, mean age = 56.7 years) and 11 healthy control participants who met these criteria (Table 1).

**Table 1.** Demographics.

| subID | Age | Gender | Education | Handedness | ETDRSSnellen |
| --- | --- | --- | --- | --- | --- |
| MDP005 | 22 | M | Associate degree | R | 125 |
| MDP006 | 50 | F | Bachelor's degree | R | 160 |
| MDP008 | 70 | F | Associate degree | L | 160 |
| MDP014 | 68 | M | Master's degree | R | 400 |
| MDP016 | 55 | M | Some college, no degree | R | 150 |
| MDP021 | 27 | M | No diploma, but finished 12th Grade | R | 200 |
| MDP022 | 83 | F | Some college, no degree | R | 100 |
| MDP023 | 35 | F | Master's degree | R | 400 |
| MDP050 | 83 | M | Doctoral degree | R | 125 |
| MDP065 | 61 | F | 12th Grade, high school graduate | R | 160 |
| MDP122 | 70 | F | Bachelor's degree | R | 400 |

All methods, including obtaining informed consent, were carried out in accordance with ethical standards under the oversight of the University of Alabama at Birmingham (UAB) Institutional Review Board.

**Figure 1.**
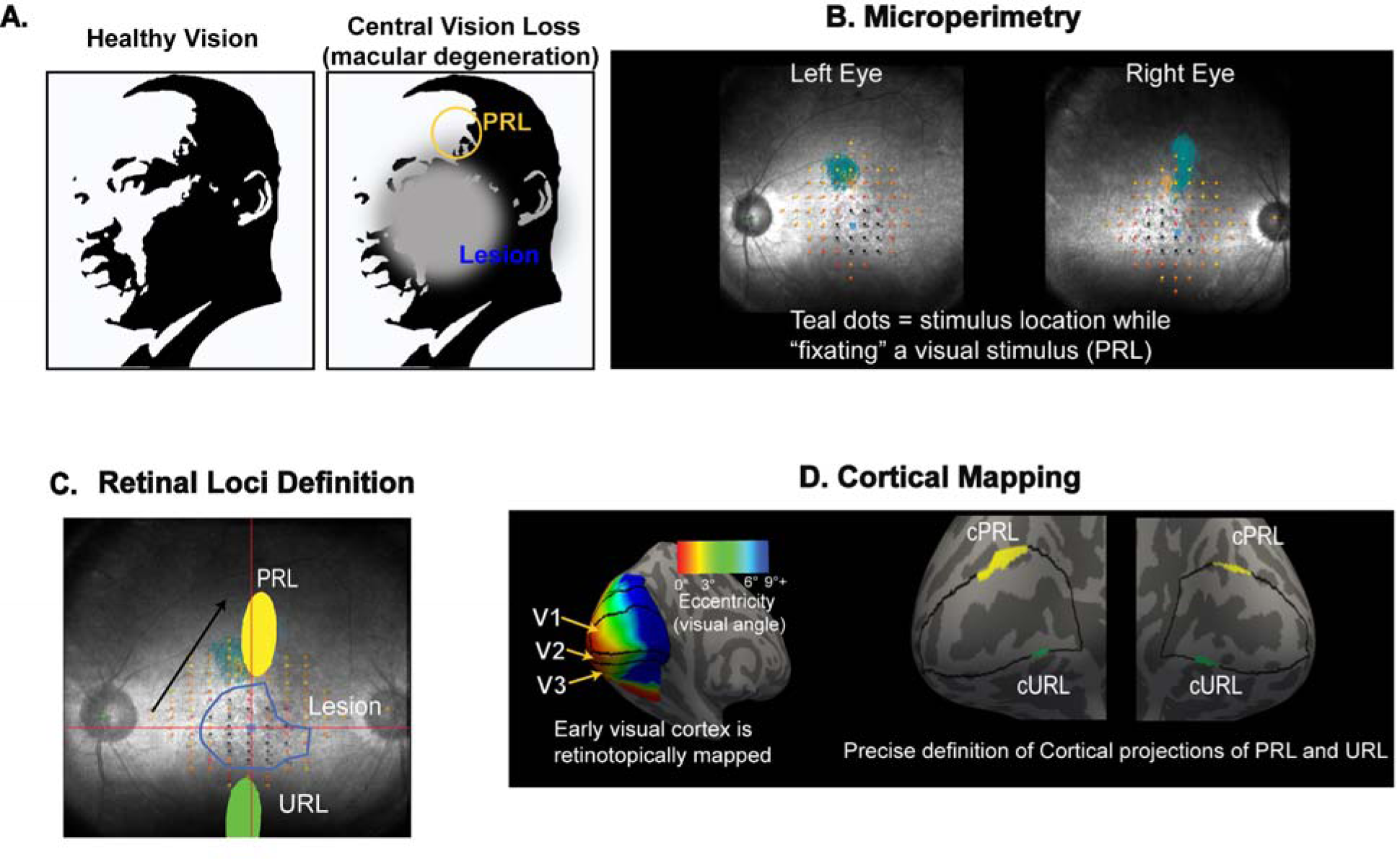
Schematic of Early Visual Cortex ROI Definition. Conceptualization of the perceptual experience of individuals with macular degeneration in comparison to healthy vision is shown in (A). In macular degeneration, a lesion forms in the center of the retina, rendering patients unable to see in the center of the visual field (gray patch). Retinal imaging was first conducted using microperimetry separately in both eyes (B). The PRL (preferred retinal locus), URL (un-preferred retinal locus), and lesion are then determined (C). Using this information, a retinotopic atlas of visual cortex is then is used (D - left) to map the cortical representations of these three loci in early visual cortex (V1, V2, V3), shown here right for V1 (D).

### MRI Data Acquisition

Functional and structural imaging data were acquired using a 3T Siemens Magnetom Prisma MRI scanner using protocols based on those of the Human Connectome Project (Glasser et al., 2016b). High-resolution 3D anatomical scans were obtained (T1-weighted; repetition time (TR) = 2400 ms; echo time (TE) = 2.22 ms; field of view (FOV(ap,rl,fh)) = 208 X 208 X 144 mm; voxel size = 0.8 mm isotropic; flip angle (FA) = 8 deg). T1w images were reacquired if the initial acquisition was of poor quality. Resting-state functional scans (eyes open) were acquired in total darkness (T2*- weighted, TR=800 ms, TE= 37 ms, FA = 52°; voxel size = 2.0 X 2.0 X 2.0 mm isotropic; echo spacing = 0.58 ms; mutli-band acceleration factor = 8) resulting in 420 volumes per scan. In order to ensure complete darkness, the investigators blocked out all possible sources of light in the room (windows, waveguides, lights on equipment) prior to the start of the resting-state scan. Participants were instructed to relax and keep their eyes open until the scan was complete. An infrared eye tracking system, which does not emit visible light, was used to monitor participants during scanning in order to ensure that their eyes remained open during the scanning session. Each scan lasted a total of 5.6 minutes and was repeated 8 times for a total of 44.8 minutes of total scan time.

### MRI Data Processing

Raw data MRI data files were first converted into Brain Imaging Data Structure (BIDS) format (Gorgolewski et al., 2016) in order to enable use with open source pre- processing pipelines. Initial first-pass quality control was performed through manual inspection of the data using MRIQC (Esteban et al., 2017). Initial image preprocessing steps were performed with FMRIPrep version 1.2.5 (https://fmriprep.org/) under the default settings (Esteban et al., 2019). For reproducibility of methods, a more detailed description is provided in the supplementary material. To summarize briefly: BOLD data underwent co-registration to T1w anatomical space, spatiotemporally filtered, slice-time corrected, scrubbed for motion artifacts using the XCP engine workflow (Ciric et al., 2018) with a framewise displacement threshold of 0.5mm. Data were then converted into fsLR_32k cifti template-space using *Ciftify (Dickie et al., 2019)* and tools from the *HCP Connectome Workbench (Marcus et al., 2011)*

### Region of Interest Definitions

Regions of interest within the early visual cortex were defined based on atlases of visual regions coupled (described fully in the next paragraph) with maps of the visual field acquired from retinal imaging acquired outside of the MRI scanner (as described in this paragraph). Visual field mapping was performed on a Centervue Macular integrity Assessment (MAIA) device (Centervue, Padova, Italy). The MAIA device uses a confocal scanning laser ophthalmoscope to image the retina in real-time while perimetry is performed using a Goldmann size III white stimulus. During the examination, the MAIA uses a 25 Hz eye tracker to compensate for eye movements and measure fixation. Participants fixate a red annulus and indicate when they detect a light flash with a button push. A 4-2 staircase strategy was used to determine sensitivity thresholds. This resulted in three pieces of information: a map of the retinal lesion, a map of the retinal locations with spared and lesioned vision, as well as an estimate of the location at which the MD patients tended to primarily fixate (i.e., the preferred retinal locus, PRL). The PRL was defined based on the cloud of fixation locations over time. The center of the PRL was defined as the center of the cloud, and the boundary was defined as the Bivariate contour ellipse area (BCEA) that included 63% of fixation locations (Steinman, 1965). In cases where the PRL was located in extremely eccentric areas, the 95% BCEA was used in order to capture enough cortical vertices for each ROI. The boundaries of the retinal lesion were manually drawn from the MAIA output image based on the locations where the participant did not detect light.

**Figure 2.**
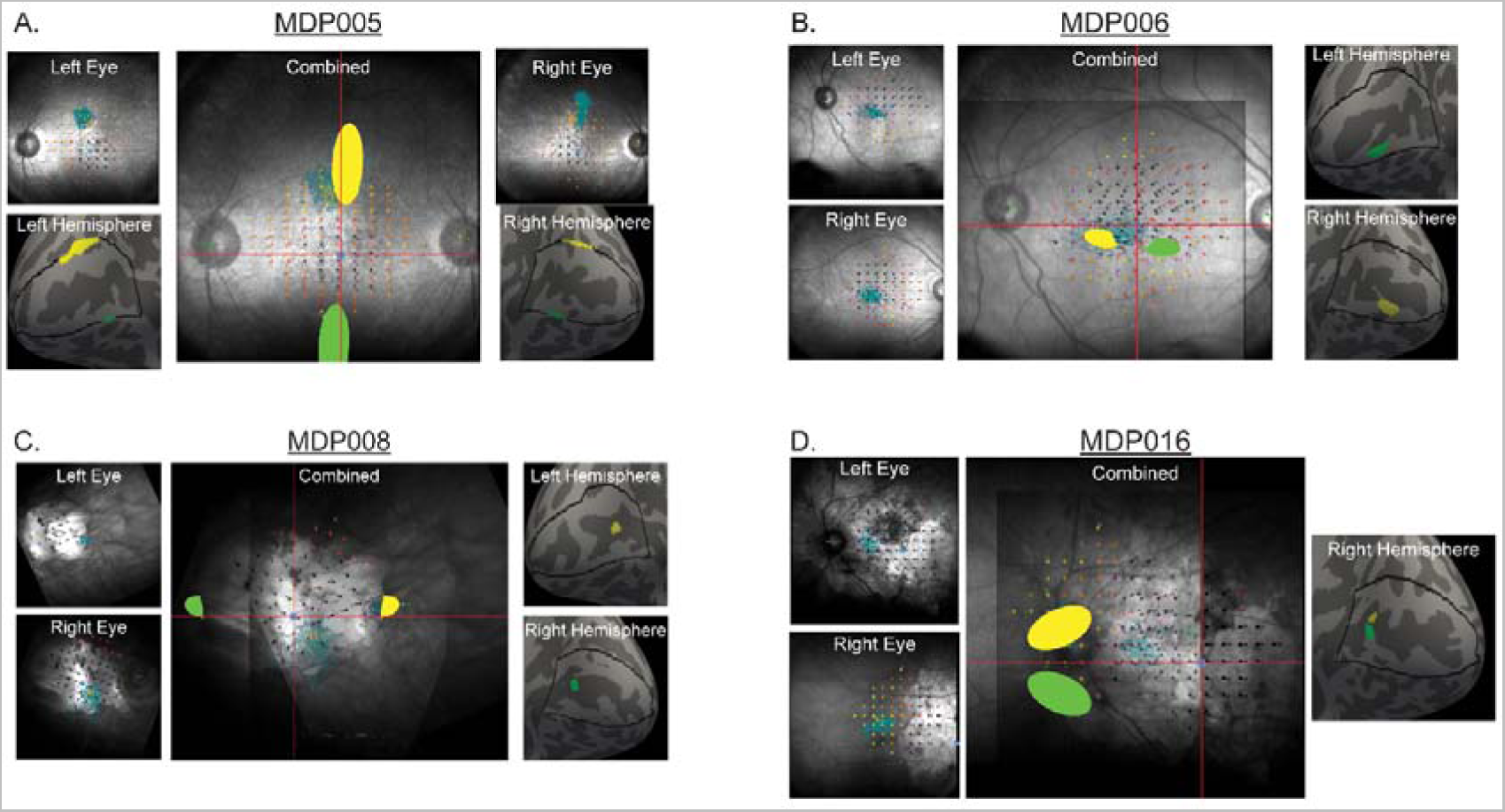
Individual Participant Retinal Images. Retinal images from microperimetry are shown for 4 representative individuals. PRL locations are shown in yellow. URL locations are shown in green. Microperimetry images for left and right eyes are shown in upper corners of each panel. Center images show combined image of left and right retinal images. Mapping onto left and right cortical surfaces are shown in bottom corners of each panel.

The locations of the PRL region and the retinal lesion were mapped to their corresponding regions on the cortical surface in V1, V2, and V3 using a retinotopic atlas (Benson and Winawer, 2018; Benson et al., 2012; Defenderfer et al., 2021). This resulted in regions of interest for each MD participant in V1, V2 and V3 for the cortical representation of the PRL (cPRL) and the cortical representation of the lesion (i.e. - the “lesion project zone”, LPZ). In addition to the cPRL and LPZ regions, we also defined a control region that we refer to here as the cortical representation of the “Un-preferred Retinal Locus” or c*URL.* This region was defined by identifying a region on the retina outside of the lesion that was: 1) the same eccentricity as the PRL region, 2) as far away from the PRL as possible (in degrees polar angle), and 3) could be confirmed to have light sensitivity as measured by the MAIA. We treated each healthy control participant the same, placing their cortical ROIs according to their matched MD participant’s visual field information from the MAIA. These regions were first created on the native cortical surface, and then projected onto the fsLR_32k template in cifti space.

In addition to the early visual cortex ROIs, we used a previously published cortical atlas (Glasser et al., 2016a) to define regions of interest for three category selective visual regions: fusiform face area, parahippocampal area, and middle temporal area.

### Functional Connectivity analysis

The average time course of each ROI was extracted from the preprocessed resting state fMRI data using in-house MATLAB scripts. Functional connectivity was then calculated as the Pearson’s correlation between the timecourses of each early visual cortex ROI and those of the category selective ROIs. Connectivity for each early visual cortex ROI (cPRL, cURL, and LPZ) was first calculated independently at the levels of V1, V2, and V3 and then averaged across levels to generate a single value for each ROI type.

### Statistical analysis

The effects of deprivation, preferred usage, and non-preferred usage on functional connectivity were assessed by using two-way, repeated measures analysis of variance (ANOVA) with factors of early visual cortex ROI (LPZ, cPRL, and cURL) and group (MD vs. healthy controls). Here, we applied a “yoked-control” approach, using group as a repeated measures factor due to the fact that each MD participant was matched based on age, gender, and education to a control participant. Each MD participant’s individualized regions of interest for LPZ, cPRL, and cURL were applied to their yoked healthy vision control participant. This approach helps control for factors of age, gender, and education. Significant results using this approach were followed up with unpaired tests as a validation measure (described in results). In cases where statistical assumptions were not met, Wilcoxon rank sum tests were used as a non-parametric alternative to the ANOVA. Statistical tests were corrected for multiple comparisons using the Bonferroni correction procedure.

## Results

### Functional Connectivity to FFA

A two-way repeated measures ANOVA revealed a statistically significant main effect of early visual cortex ROI (LPZ, cPRL, cURL) on functional connectivity to fusiform face area (F(2,20) = 78.712, p = 3.31e-10). Post-hoc t-tests with a Bonferroni adjustment revealed that functional connectivity of the LPZ was significantly greater than that of the cPRL and cURL in both MD patients and healthy controls (Fig. 3). No significant main effect of group was observed, nor was there a significant interaction between group and early visual cortex ROI.

**Figure 3.**
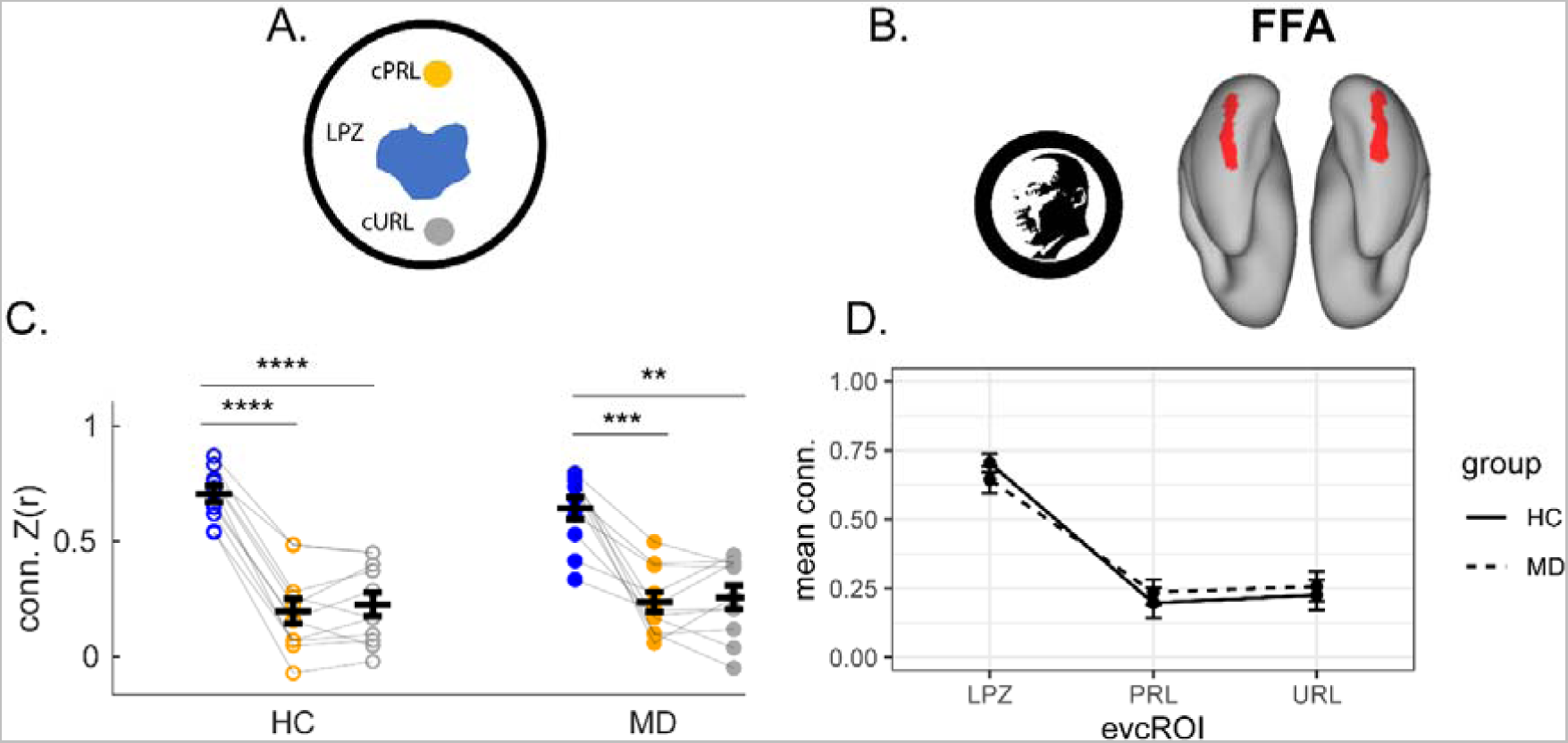
Connectivity to Fusiform Face Area. Functional connectivity to fusiform face area (B) was measured for the LPZ (blue), the cPRL (yellow), and the cURL (gray). Connectivity to FFA was significantly higher for the LPZ, relative to the cPRL and cURL, in both groups (C). Group means for each ROI are shown in D. Error bars in both figures represent the standard error of the mean.

### Functional Connectivity to PHA

We first tested whether central vision loss and increased use of the PRL is related to altered patterns of functional connectivity between early visual cortex and parahippocampal area, a brain region that demonstrates selectivity for scenes. To this, we conducted a two-way, repeated measures ANOVAs with factors of group (MD vs. healthy controls, as a yoked control) and early visual cortex ROI (LPZ, PRL, and URL). The ANOVA results revealed a statistically significant main effect of early visual cortex ROI (LPZ, PRL, URL) on functional connectivity to parahippocampal area (F(2,20) = 32.70, p = 4.96e-7) Post-hoc t-tests with a Bonferroni adjustment revealed that functional connectivity of the LPZ was significantly greater than that of the PRL and URL in both MD patients and healthy controls. We did not observe a significant main effect of group, nor was there a significant interaction between group and ROI.

**Figure 4.**
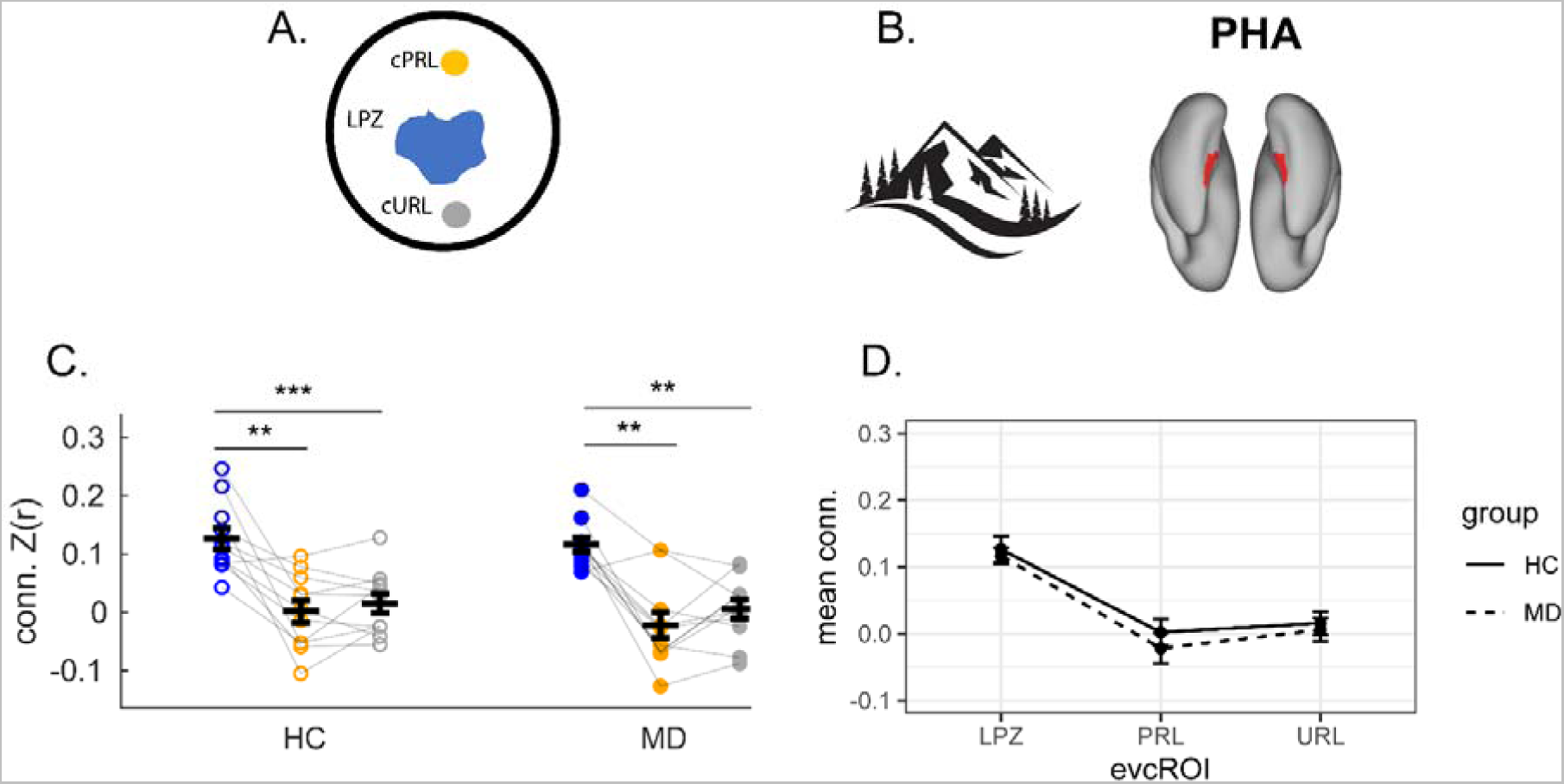
Connectivity to Parahippocampal Area. Functional connectivity was measured between three region of early visual cortex (A): LPZ (blue), the cPRL (yellow), and the cURL (gray) and parahippocampal area (B). A two-way, repeated-measures ANOVA revealed a significant main effect of early visual cortex ROI, such that LPZ connectiviy was higher relative to both the cPRL and cURL in both groups (C). Group means for each ROI are shown in D. Error bars in both figures represent the standard error of the mean.

### Functional Connectivity to MT

A Levene’s test of homogeneity of variances test showed unequal variances connectivity between groups for the peripheral ROIs (cPRL: p=.01, cURL: p=.01). Specifically, MD participants are more homogeneous than controls. Because there is no well-accepted alternative for 2-way ANOVA under unequal variances, and because we did not see differences between the connectivity of the cPRL and cURL, we followed up with a nonparametric Wilcoxon test comparing MD participants to controls. To this end we calculated a mean score for the central ROI (LPZ) and subtracted this value from the mean of the peripheral rois (cPRL and cURL). This “peripheral relative to central” value was then compared between the two groups. We found that peripheral relative to central connectivity to MT was significantly different between groups using both a paired/yoked approach, where each MD participant matched by age, gender, and education to a control ( p=4.88e-3) and an unpaired approach (p=1.40e-2).

In healthy controls, peripheral minus central connectivity values were centered around zero (Figure 5E), suggesting very little difference between the ROIs. In MD, this number was a positive value, suggesting that peripheral connectivity to MT was greater than that of central representations (mean = 0.12).

**Figure 5.**
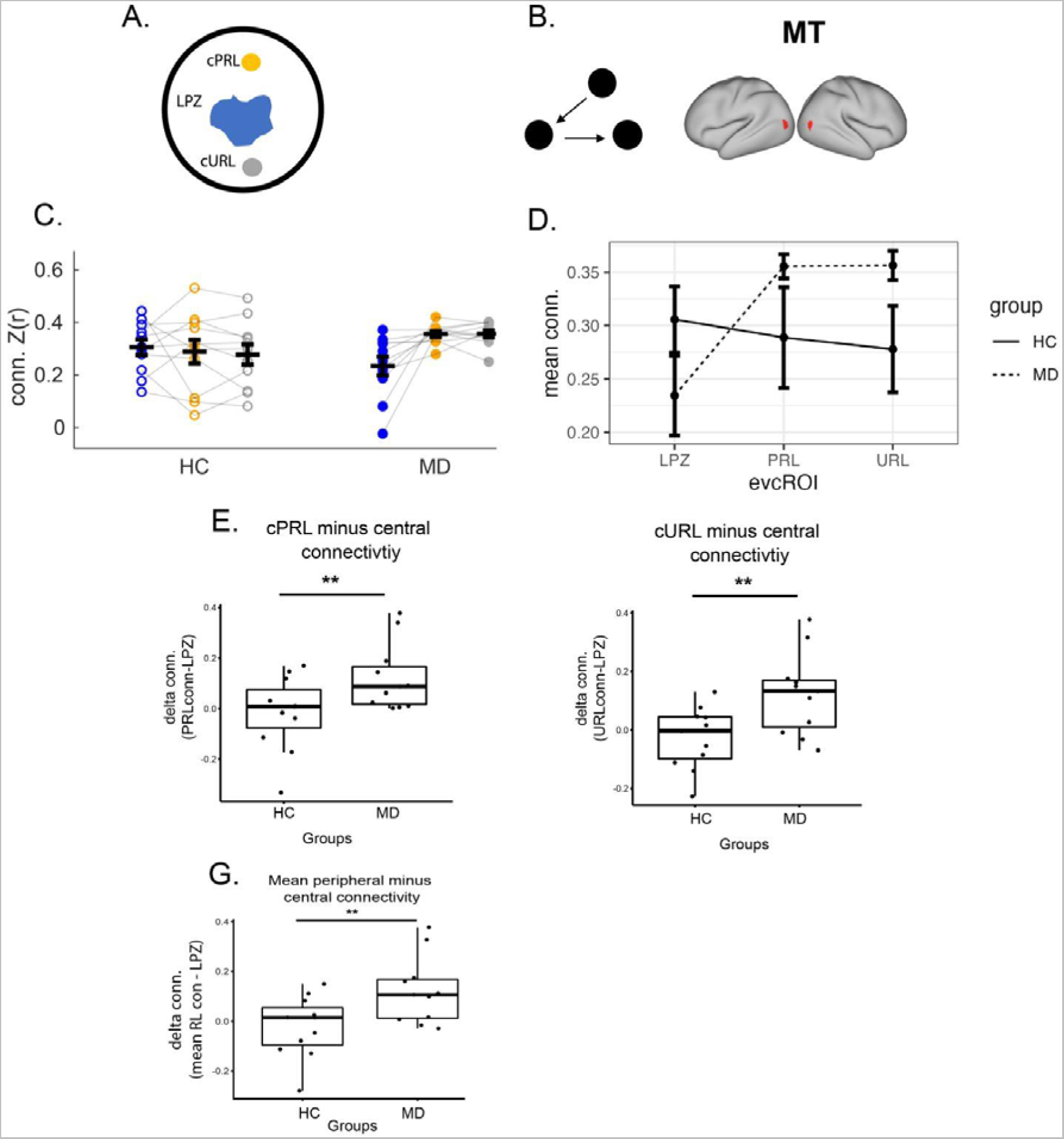
Connectivity to Middle Temporal Area. Functional connectivity was measured between three regions of Early Visual Cortex (A): LPZ (blue), the cPRL (yellow), and the cURL (gray) and MT (B). Connectivity between each early visual cortex ROI and MT are shown in for visualization purposes only (C). Groups means are shown in (D). Wilcoxon rank sum tests revealed that cPRL connectivity (E) and cURL connectivity (F) relative to central (LPZ) connectivity were significantly greater in MD participants relative to healthy vision controls (both p<0.01). Similar results were found when examining the mean of the two peripheral ROIs (cPRL and cURL) relative to central (LPZ) connectivity (G).

### Generalizability of Peripheral Connectivity

The highly similar functional connectivity profiles for the cPRL and the cURL regions suggested the observed increase in peripheral connectivity may not be specific to only the preferentially used portion of peripheral vision. As a result, we tested whether a similar result would be found if we looked at all areas of usable peripheral vision simultaneously. To do this, we created ROIs for each individual subject pair that included all areas of usable vision outside of the lesion projection zone. A Levene’s test of homogeneity of variances test showed unequal variances connectivity between groups for the peripheral ROIs (cPRL: p=.01, cURL: p=.01). Specifically MD participants were more homogeneous than controls. Similar to described earlier, an “all peripheral relative to central” score was calculated by subtracting LPZ to MT connectivity from “all peripheral” connectivity. A Wilcoxon rank sum test revealed that peripheral (relative to central) connectivity was significantly greater in the MD group compared to controls using both a paired (p=6.84e-3) and unpaired approach (p=3.80e-2).

**Figure 6.**
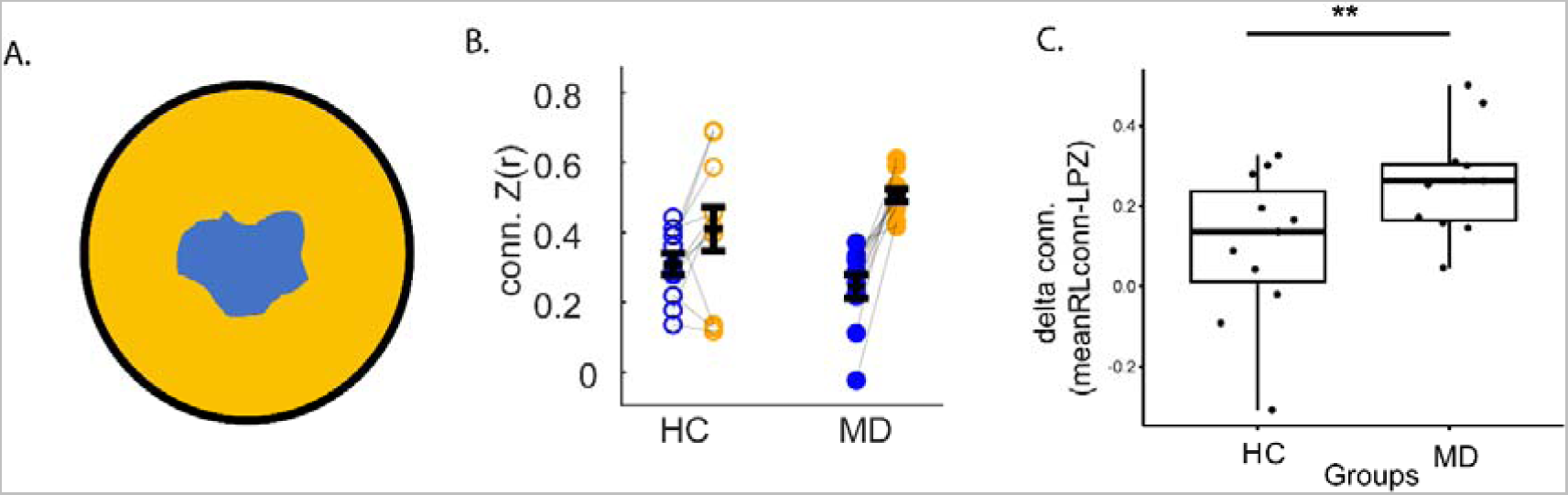
Generalizability of peripheral connectivity difference. Connectivity of all peripheral regions to MT was calculated in order to probe the generalizability of the previous observed effects. Schematic of the all peripheral regions (yellow) and lesion region (blue) are shown (A). Connectivity between each early visual cortex ROI and MT are shown in (B). A direct comparison of “all peripheral” minus LPZ difference scores are also shown (C), revealing greater peripheral relative to central connectivity in MD participants (p<.0.01). Statistical analysis in (C) was performed using a Wilcoxon rank sum test. Error bars for all figures represent the standard error of the mean.

## Discussion

We explored how three key features of sensory experience (deprivation, maintenance, and preferential usage) influence the functional connection patterns of the visual cortex. Our findings, using central vision loss as a model, confirm prior evidence demonstrating that early visual cortex exhibits retinotopically specific patterns of functional connectivity to downstream, category-selective visual areas. Additionally, we show that central vision loss is associated with strengthened functional connectivity between cortical representations of peripheral vision and regions involved in motion detection (MT). Interestingly, we found that these changes were present not just for the region of preferred usage, but extended to other regions of usable peripheral vision. Together, these findings suggest that long-term changes in visual experience can produce changes in intrinsic cortical activity, well after the critical period of visual development.

### Retinotopic patterns of connectivity to FFA and PHA

Functional connections between early visual cortex and category-selective visual areas follow retinotopically specific patterns. We observed that compared to peripheral representations in early visual cortex, centrally representing regions exhibited greater functional connectivity to both FFA and PHA. The stronger connectivity of FFA to central representations is consistent with previous work showing that activity in FFA is biased towards information from central vision (Levy et al., 2001); (Hasson et al., 2002). Here, we replicate this finding by showing in MD patients and healthy controls that the cortical representation of central vision in early visual cortex (the LPZ) was more strongly connected to FFA than peripheral regions (cPRL and cURL). This bias towards central vision information is believed to be due to the high visual acuity required to make out details in faces that convey important information, such as facial expressions, physical facial features, and eye gaze direction (Levy et al., 2001). Because individuals with central vision loss must rely on peripheral vision to make out facial features, we initially hypothesized that FFA would exhibit a bias towards peripheral visual information in patients with MD. However, we did not find a statistically significant difference in FFA connectivity between groups. There was considerable between-subject variability, however, and future work will be needed to identify whether this variability may be explained by other factors. An important question is whether brain connectivity, along with the decreased visual acuity inherent in central vision loss, contributes to the known deficits that MD patients commonly experience difficulty with tasks like recognizing faces (Bullimore et al., 1991). It may be possible that without strengthened peripheral connectivity, information from the peripheral visual field is not properly transmitted to regions like FFA in a way that would improve visual performance on facial processing tasks. Future studies should explore this question and determine whether the strength of connectivity between FFA and peripheral representation in early visual cortex relate to performance on peripherally presented facial stimuli, and whether this connectivity can be strengthened with intervention.

Another possible reason for the lack of stronger connectivity between cPRL and FFA in patients with MD may be that individuals with central vision loss may be using vision less overall for visual tasks (like facial recognition) that require fine-grain detail. The low resolution of patients’ spared vision may not allow individuals to engage in enough everyday facial recognition to produce noticeable changes in connectivity. In future work, it will be important to factor in whether the amount of overall vision use in MD patients is related to the strength of connectivity between peripheral representations and regions like FFA, that are involved in tasks normally performed with central vision.

We found that PHA showed stronger connectivity to the representations of central vision (LPZ) than peripheral vision in both MD patients and controls. This finding was not in the direction of previous work showing PHA may exhibit bias towards peripheral visual information (Baldassano et al., 2016; Levy et al., 2001). However, it should be noted that different subregions of PHA have been shown to have differential patterns of connection to V1 (Baldassano et al., 2016) and our PHA region was defined as a relatively large swath of cortex, and not defined based on localizers as in some earlier studies. Here, a larger region of interest was used for PHA in an attempt to account for previous reports indicating its high individual variability in location and size (Zhen et al., 2017). Future work is needed to probe whether these more subtle relationships between subregions within the parahippocampal area and early visual cortex change following the loss of central vision.

### Functional connectivity to MT

In participants with healthy vision, functional connections between MT and early visual cortex did not significantly differ between central representations (LPZ) and peripheral representations (cPRL and cURL). Patients with central vision loss, but not controls, showed retinotopic patterns of connectivity: In patients, there was stronger functional connectivity between MT and representations of peripheral vision than central vision. Importantly, this between-group difference was statistically significant, indicating that the presence of the retinotopic pattern may be a consistent outcome of the experience of central vision loss. Notably, the relative increase in functional connectivity for peripheral vision was not confined to regions within the vicinity of the cPRL and cURL, but extended to the cortical representation of all of the visual periphery.

Our finding of increased connectivity between peripheral representations in early visual cortex and area MT is consistent with previous reports in the literature. For example, prior work in animal models of MD show that central vision loss is associated with improved motion detection as well as velocity discrimination (Burnat et al., 2017). Other work has demonstrated that connections between MT and early visual cortex are modifiable in adulthood and can boost sensitivity to motion (Romei et al., 2016). This raises the question of whether the greater connectivity between peripheral representations and MT in our MD patients is associated with enhanced performance on motion detection tasks. In everyday life, the ability to detect motion is commonly used for visual tasks like spatial navigation (Wolbers et al., 2007). For example, walking through a house involves interacting with many “looming” stimuli such as doorways, whose projection on the retina moves quickly in the periphery. In contrast to tasks like reading or recognizing faces, which are normally performed with foveal vision, activities like spatial navigation rarely rely on fine detail. For this reason, MD patients are likely to regularly engage in spatial navigation involving peripheral looming stimuli, but do less face recognition. Thus, repeated use of spared peripheral vision in MD patients for an activity like spatial navigation may selectively lead to changes in functional connectivity to areas like MT, which are essential for moving through the world. Future work should consider whether the functional connectivity of MT is related to behavioral performance on motion tasks in MD patients.

Previous work has shown that while other brain regions in the visual network decrease their overall activity level, activity in MT is still maintained after central retinal lesioning (Burnat et al., 2017), despite the loss of bottom up inputs from central vision. This suggests that MT’s increased connectivity to peripheral representations in early visual cortex in our data may act to maintain a homeostatic level of activity in MT. However, future studies would need to confirm this notion by looking at the overall activity level of MT (using methods like positron emission tomography) in relation to increased functional connectivity to peripheral regions in early visual cortex.

Understanding the inputs to MT can give insight into why MT connectivity to peripheral representations in early visual cortex increases in patients with central vision loss. The majority of inputs to MT from V1 come from the magnocellular stream, which is known to have cells with relatively large receptive fields and strong sensitivity to motion. However, MT also contains some inputs from the parvocellular stream, which dominate the makeup of ganglion cells in the central retina (Dacey, 1993). Removal of input from the fovea, which is dominated by less motion-sensitive parvocellular neurons (Masri et al., 2020; Yan et al., 2020), may result in a proportional increase of input to V1 from motion sensitive cells (magnocellular) compared to non-motion sensitive (parvocellular) cells. Thus, removal of parvocellular input may allow V1 to be more strongly driven by magnocellular inputs, resulting in overall greater functional connectivity of peripheral regions to MT.

### Generalizability of peripheral connectivity effects

Interestingly, we found that the effect of increased connectivity between MT and peripheral early visual cortex was present not only for the area of preferred usage (PRL), but also for un-preferred areas. Furthermore, this effect was maintained when factoring in the cortical representation of all parts of usable peripheral vision. This finding seems logical within the grand scope of PRL usage in MD patients. Prior reports indicate that individuals with MD often use multiple PRLs for different tasks (Déruaz et al., 2002; Duret et al., 1999; Lei and Schuchard, 1997). In our study, PRLs were defined based on microperimetry in each participant’s better eye during performance of only one task. Thus, our “unpreferred retinal locations” may include some locations that are preferred during different tasks, and the generalization of the connectivity effects may result from this effect. On the other hand, the connectivity difference was shown to generalize to the cortical representation of the entire spared peripheral visual field. This would not be expected if the increased connectivity were limited to a handful of specific retinal locations. Thus, our data suggest that MT increases connection strength to the cortical representations of all spared vision after central vision loss, regardless of preferential use. This interpretation is consistent with the idea that habitual processing of large peripheral looming stimuli are involved in this shift in connection strength. However, more work is needed to clarify this relationship.

### Differences in variability of functional connections to MT

In this analysis, we detected group differences in variability for functional connectivity between peripheral representations in early visual cortex and MT. While comparisons of variability are often used as a measure to test assumptions for statistical analysis, in the case of plasticity, individual variability in functional connectivity can be informative of deeper underlying biology (Mueller et al., 2013). Furthermore, individual variability has been shown to be important in several contexts related to normal development, aging, and brain-related diseases (Hahamy et al., 2015; Sele et al., 2021). Notably, previous work suggests that group differences in variability exist when examining individuals with healthy vision vs. individuals with congenital blindness. For example, Sen and colleagues (Sen et al., 2022) found that individual variability in the functional connectivity of visual cortex is higher in individuals with congenital blindness compared to individuals with healthy vision. These results suggest that increased use of vision results in more consistent functional connectivity between individuals. In our study, we observed that increased use of peripheral vision similarly leads to more across-subject similarity in connectivity in individuals with MD. This serves as yet another form of evidence toward the importance of visual experience in the establishment of connectivity between brain regions. Moreover, it supports the idea put forth by Sen and colleagues that “shared sensory experience enforces consistency across individuals.” Our results add a new layer to this idea, suggesting that visual experience can influence individual variability in a specific manner when one particular part of the visual field is used more. While these differences in individual variability are interesting, a couple of outstanding questions remain. First, the results shown in the current study only look at individuals who had visual input in the early stages of life.

Previous work has shown visual system development in sighted individuals decreases variability compared to congenitally blind individuals (Sen et al., 2022). Our results suggest a possible further refinement of this individual variability in adulthood when individuals increasingly use one section of the visual field. Second, it remains to be fully understood why increased use is related to reduced variability of functional connectivity in visual cortex. One possibility may be activity-dependent changes that are known to drive plasticity related-change in visual cortex (Hubel and Wiesel, 1963; Hubel and Wiesel, 1970). In the case of central vision loss, increased reliance on peripheral vision likely drives higher-than-normal levels of activity that may alter peripheral representations of visual cortex that look similar across individuals. Such activity may drive activity-dependent gene-expression changes that have been previously shown to play an important role in the plasticity of visual cortex (Tropea et al., 2006). In the present study, our findings highlight an important role of experience in shaping individual differences in the brain that may be related to such changes.

### Conclusions

In summary, our study sheds light on the impact of preferential usage on patterns of functional connections, specifically in individuals with long-term loss of central vision. Our findings provide strong evidence for the role of brain plasticity well after the critical period of development, demonstrating Hebbian-like plasticity in the connections between early visual cortex and the middle temporal area. Moreover, our results highlight the importance of functional connectivity during rest as a meaningful characteristic of the brain that can be shaped by experience. Overall, this study highlights the remarkable flexibility of the human brain and its capacity for adaptation in response to changes in sensory input, even in older adults with long-term visual impairment. These findings provide new insights into the mechanisms of brain plasticity and have important implications for the development of new approaches for rehabilitation and treatment.

## Acknowledgments

The authors would like to acknowledge The UAB Callahan Clinical Research Unit, Dr. Cynthia Owsley, Mark Clark, and the Civilian International Research Center, as well as our funding sources:

## Author contributions

L.L.F - conceptualization, formal analysis, investigation, writing - original draft; M.D. - methodology, investigation; P.D. - methodology, investigation; P.S. - software, formal analysis; D.K.D. - resources, supervision; K.MV. - conceptualization, writing -review and editing, funding acquisition, supervision

## Data availability statement

The data associated with this project are currently available as part of the Human Connectome Project: Connectomes in Diseases repository.

## Ethics approval statement

All portions of this study were carried out in accordance with ethical standards under the oversight of the University of Alabama at Birmingham (UAB) Institutional Review Board.

## Patient consent statement

Informed consent was obtained from all participants prior to study enrollment.

## Permission to reproduce material from other sources

n/a

## Clinical trial registration

n/a

## Bibliography

Andrews-Hanna JR, Snyder AZ, Vincent JL, Lustig C, Head D, Raichle ME, Buckner RL (2007): Disruption of large-scale brain systems in advanced aging. Neuron 56:924– 935.

Arcaro MJ, Honey CJ, Mruczek REB, Kastner S, Hasson U (2015): Widespread correlation patterns of fMRI signal across visual cortex reflect eccentricity organization. eLife 4.

Baker CI, Dilks DD, Peli E, Kanwisher N (2008): Reorganization of visual processing in macular degeneration: replication and clues about the role of foveal loss. Vision Res 48:1910–1919.

Baker CI, Peli E, Knouf N, Kanwisher NG (2005): Reorganization of visual processing in macular degeneration. J Neurosci 25:614–618.

Baldassano C, Fei-Fei L, Beck DM (2016): Pinpointing the peripheral bias in neural scene-processing networks during natural viewing. J Vis 16:9.

Baseler HA, Gouws A, Haak KV, Racey C, Crossland MD, Tufail A, Rubin GS, Cornelissen FW, Morland AB (2011): Large-scale remapping of visual cortex is absent in adult humans with macular degeneration. Nat Neurosci 14:649–655.

Benson NC, Butt OH, Datta R, Radoeva PD, Brainard DH, Aguirre GK (2012): The retinotopic organization of striate cortex is well predicted by surface topology. Curr Biol 22:2081–2085.

Benson NC, Winawer J (2018): Bayesian analysis of retinotopic maps. eLife 7.

Biswal B, Yetkin FZ, Haughton VM, Hyde JS (1995): Functional connectivity in the motor cortex of resting human brain using echo-planar MRI. Magn Reson Med 34:537–541.

Bullimore MA, Bailey IL, Wacker RT (1991): Face recognition in age-related maculopathy. Invest Ophthalmol Vis Sci 32:2020–2029.

Burge WK, Griffis JC, Nenert R, Elkhetali A, DeCarlo DK, ver Hoef LW, Ross LA, Visscher KM (2016): Cortical thickness in human V1 associated with central vision loss. Sci Rep 6:23268.

Burnat K, Hu T-T, Kossut M, Eysel UT, Arckens L (2017): Plasticity Beyond V1: Reinforcement of Motion Perception upon Binocular Central Retinal Lesions in Adulthood. J Neurosci 37:8989–8999.

Calford MB, Wang C, Taglianetti V, Waleszczyk WJ, Burke W, Dreher B (2000): Plasticity in adult cat visual cortex (area 17) following circumscribed monocular lesions of all retinal layers. J Physiol (Lond) 524 Pt 2:587–602.

Chang EF, Merzenich MM (2003): Environmental noise retards auditory cortical development. Science 300:498–502.

Chan MY, Park DC, Savalia NK, Petersen SE, Wig GS (2014): Decreased segregation of brain systems across the healthy adult lifespan. Proc Natl Acad Sci USA 111:E4997–5006.

Ciric R, Rosen AFG, Erus G, Cieslak M, Adebimpe A, Cook PA, Bassett DS, Davatzikos C, Wolf DH, Satterthwaite TD (2018): Mitigating head motion artifact in functional connectivity MRI. Nat Protoc 13:2801–2826.

Cramer SC, Sur M, Dobkin BH, O’Brien C, Sanger TD, Trojanowski JQ, Rumsey JM, Hicks R, Cameron J, Chen D, Chen WG, Cohen LG, deCharms C, Duffy CJ, Eden GF, Fetz EE, Filart R, Freund M, Grant SJ, Haber S, Vinogradov S (2011): Harnessing neuroplasticity for clinical applications. Brain 134:1591–1609.

Dacey DM (1993): The mosaic of midget ganglion cells in the human retina. J Neurosci 13:5334–5355.

Defenderfer M, Demirayak P, Visscher KM (2021): A method for mapping retinal images in early visual cortical areas. Neuroimage 245:118737.

Déruaz A, Whatham AR, Mermoud C, Safran AB (2002): Reading with multiple preferred retinal loci: implications for training a more efficient reading strategy. Vision Res 42:2947–2957.

Dickie EW, Anticevic A, Smith DE, Coalson TS, Manogaran M, Calarco N, Viviano JD, Glasser MF, Van Essen DC, Voineskos AN (2019): Ciftify: A framework for surface-based analysis of legacy MR acquisitions. Neuroimage 197:818–826.

Dilks DD, Baker CI, Peli E, Kanwisher N (2009): Reorganization of visual processing in macular degeneration is not specific to the “preferred retinal locus”. J Neurosci 29:2768–2773.

Dosenbach NUF, Nardos B, Cohen AL, Fair DA, Power JD, Church JA, Nelson SM, Wig GS, Vogel AC, Lessov-Schlaggar CN, Barnes KA, Dubis JW, Feczko E, Coalson RS, Pruett JR, Barch DM, Petersen SE, Schlaggar BL (2010): Prediction of individual brain maturity using fMRI. Science 329:1358–1361.

Duret F, Issenhuth M, Safran AB (1999): Combined use of several preferred retinal loci in patients with macular disorders when reading single words. Vision Res 39:873– 879.

Esteban O, Birman D, Schaer M, Koyejo OO, Poldrack RA, Gorgolewski KJ (2017): MRIQC: Advancing the automatic prediction of image quality in MRI from unseen sites. PLoS ONE 12:e0184661.

Esteban O, Markiewicz CJ, Blair RW, Moodie CA, Isik AI, Erramuzpe A, Kent JD, Goncalves M, DuPre E, Snyder M, Oya H, Ghosh SS, Wright J, Durnez J, Poldrack RA, Gorgolewski KJ (2019): fMRIPrep: a robust preprocessing pipeline for functional MRI. Nat Methods 16:111–116.

Fair DA, Dosenbach NUF, Church JA, Cohen AL, Brahmbhatt S, Miezin FM, Barch DM, Raichle ME, Petersen SE, Schlaggar BL (2007): Development of distinct control networks through segregation and integration. Proc Natl Acad Sci USA 104:13507– 13512.

Ferris FL, Kassoff A, Bresnick GH, Bailey I (1982): New visual acuity charts for clinical research. Am J Ophthalmol 94:91–96.

Fox MD, Raichle ME (2007): Spontaneous fluctuations in brain activity observed with functional magnetic resonance imaging. Nat Rev Neurosci 8:700–711.

Geerligs L, Renken RJ, Saliasi E, Maurits NM, Lorist MM (2015): A Brain-Wide Study of Age-Related Changes in Functional Connectivity. Cereb Cortex 25:1987–1999.

Genç E, Schölvinck ML, Bergmann J, Singer W, Kohler A (2016): Functional connectivity patterns of visual cortex reflect its anatomical organization. Cereb Cortex 26:3719–3731.

Gilbert CD, Li W (2012): Adult visual cortical plasticity. Neuron 75:250–264.

Gilbert CD, Wiesel TN (1992): Receptive field dynamics in adult primary visual cortex. Nature 356:150–152.

Glasser MF, Coalson TS, Robinson EC, Hacker CD, Harwell J, Yacoub E, Ugurbil K, Andersson J, Beckmann CF, Jenkinson M, Smith SM, Van Essen DC (2016a): A multi-modal parcellation of human cerebral cortex. Nature 536:171–178.

Glasser MF, Smith SM, Marcus DS, Andersson JLR, Auerbach EJ, Behrens TEJ, Coalson TS, Harms MP, Jenkinson M, Moeller S, Robinson EC, Sotiropoulos SN, Xu J, Yacoub E, Ugurbil K, Van Essen DC (2016b): The Human Connectome Project’s neuroimaging approach. Nat Neurosci 19:1175–1187.

Gorgolewski KJ, Auer T, Calhoun VD, Craddock RC, Das S, Duff EP, Flandin G, Ghosh SS, Glatard T, Halchenko YO, Handwerker DA, Hanke M, Keator D, Li X, Michael Z, Maumet C, Nichols BN, Nichols TE, Pellman J, Poline J-B, Poldrack RA (2016): The brain imaging data structure, a format for organizing and describing outputs of neuroimaging experiments. Sci Data 3:160044.

Gratton C, Laumann TO, Nielsen AN, Greene DJ, Gordon EM, Gilmore AW, Nelson SM, Coalson RS, Snyder AZ, Schlaggar BL, Dosenbach NUF, Petersen SE (2018): Functional brain networks are dominated by stable group and individual factors, not cognitive or daily variation. Neuron 98:439–452.e5.

Griffis JC, Elkhetali AS, Burge WK, Chen RH, Bowman AD, Szaflarski JP, Visscher KM (2017): Retinotopic patterns of functional connectivity between V1 and large-scale brain networks during resting fixation. Neuroimage 146:1071–1083.

Grubb MS, Thompson ID (2004): The influence of early experience on the development of sensory systems. Curr Opin Neurobiol 14:503–512.

Hahamy A, Behrmann M, Malach R (2015): The idiosyncratic brain: distortion of spontaneous connectivity patterns in autism spectrum disorder. Nat Neurosci 18:302–309.

Harmelech T, Malach R (2013): Neurocognitive biases and the patterns of spontaneous correlations in the human cortex. Trends Cogn Sci (Regul Ed) 17:606–615.

Hasson U, Levy I, Behrmann M, Hendler T, Malach R (2002): Eccentricity bias as an organizing principle for human high-order object areas. Neuron 34:479–490.

Hebb D (1949): 0.(1949) The organization of behavior.

Heinzle J, Kahnt T, Haynes J-D (2011): Topographically specific functional connectivity between visual field maps in the human brain. Neuroimage 56:1426–1436.

Hubel DH, Wiesel TN (1962): Receptive fields, binocular interaction and functional architecture in the cat’s visual cortex. J Physiol (Lond) 160:106–154.

Hubel DH, Wiesel TN (1963): RECEPTIVE FIELDS OF CELLS IN STRIATE CORTEX OF VERY YOUNG, VISUALLY INEXPERIENCED KITTENS. J Neurophysiol 26:994–1002.

Hubel DH, Wiesel TN (1970): The period of susceptibility to the physiological effects of unilateral eye closure in kittens. J Physiol (Lond) 206:419–436.

James W (1890): The Principles of Psychology, vol. 1. New York (1890). Google Scholar| Crossref 1.

Kaas JH, Krubitzer LA, Chino YM, Langston AL, Polley EH, Blair N (1990): Reorganization of retinotopic cortical maps in adult mammals after lesions of the retina. Science 248:229–231.

Lei H, Schuchard RA (1997): Using two preferred retinal loci for different lighting conditions in patients with central scotomas. Invest Ophthalmol Vis Sci 38:1812– 1818.

Levy I, Hasson U, Avidan G, Hendler T, Malach R (2001): Center-periphery organization of human object areas. Nat Neurosci 4:533–539.

Lewis CM, Baldassarre A, Committeri G, Romani GL, Corbetta M (2009): Learning sculpts the spontaneous activity of the resting human brain. Proc Natl Acad Sci USA 106:17558–17563.

Liu T, Cheung S-H, Schuchard RA, Glielmi CB, Hu X, He S, Legge GE (2010): Incomplete cortical reorganization in macular degeneration. Invest Ophthalmol Vis Sci 51:6826–6834.

Marcus DS, Harwell J, Olsen T, Hodge M, Glasser MF, Prior F, Jenkinson M, Laumann T, Curtiss SW, Van Essen DC (2011): Informatics and data mining tools and strategies for the human connectome project. Front Neuroinformatics 5:4.

Masri RA, Grünert U, Martin PR (2020): Analysis of parvocellular and magnocellular visual pathways in human retina. J Neurosci 40:8132–8148.

Masuda Y, Dumoulin SO, Nakadomari S, Wandell BA (2008): V1 projection zone signals in human macular degeneration depend on task, not stimulus. Cereb Cortex 18:2483–2493.

Masuda Y, Takemura H, Terao M, Miyazaki A, Ogawa S, Horiguchi H, Nakadomari S, Matsumoto K, Nakano T, Wandell BA, Amano K (2021): V1 projection zone signals in human macular degeneration depend on task despite absence of visual stimulus. Curr Biol 31:406–412.e3.

Meunier D, Achard S, Morcom A, Bullmore E (2009): Age-related changes in modular organization of human brain functional networks. Neuroimage 44:715–723.

Mueller S, Wang D, Fox MD, Yeo BTT, Sepulcre J, Sabuncu MR, Shafee R, Lu J, Liu H (2013): Individual variability in functional connectivity architecture of the human brain. Neuron 77:586–595.

Park B-Y, Tark K-J, Shim WM, Park H (2018): Functional connectivity based parcellation of early visual cortices. Hum Brain Mapp 39:1380–1390.

Plank T, Frolo J, Brandl-Rühle S, Renner AB, Jägle H, Greenlee MW (2017): fMRI with Central Vision Loss: Effects of Fixation Locus and Stimulus Type. Optom Vis Sci 94:297–310.

Raemaekers M, Schellekens W, van Wezel RJA, Petridou N, Kristo G, Ramsey NF (2014): Patterns of resting state connectivity in human primary visual cortical areas: a 7T fMRI study. Neuroimage 84:911–921.

Raichle ME, MacLeod AM, Snyder AZ, Powers WJ, Gusnard DA, Shulman GL (2001): A default mode of brain function. Proc Natl Acad Sci USA 98:676–682.

Romei V, Chiappini E, Hibbard PB, Avenanti A (2016): Empowering Reentrant Projections from V5 to V1 Boosts Sensitivity to Motion. Curr Biol 26:2155–2160.

Sabbah N, Sanda N, Authié CN, Mohand-Saïd S, Sahel J-A, Habas C, Amedi A, Safran AB (2017): Reorganization of early visual cortex functional connectivity following selective peripheral and central visual loss. Sci Rep 7:43223.

Satterthwaite TD, Wolf DH, Erus G, Ruparel K, Elliott MA, Gennatas ED, Hopson R, Jackson C, Prabhakaran K, Bilker WB, Calkins ME, Loughead J, Smith A, Roalf DR, Hakonarson H, Verma R, Davatzikos C, Gur RC, Gur RE (2013): Functional maturation of the executive system during adolescence. J Neurosci 33:16249– 16261.

Sele S, Liem F, Mérillat S, Jäncke L (2021): Age-related decline in the brain: a longitudinal study on inter-individual variability of cortical thickness, area, volume, and cognition. Neuroimage 240:118370.

Sen S, Khalsa NN, Tong N, Ovadia-Caro S, Wang X, Bi Y, Striem-Amit E (2022): The role of visual experience in individual differences of brain connectivity. J Neurosci 42:5070–5084.

Steinman RM (1965): Effect of target size, luminance, and color on monocular fixation*. J Opt Soc Am 55:1158.

Striem-Amit E, Ovadia-Caro S, Caramazza A, Margulies DS, Villringer A, Amedi A (2015): Functional connectivity of visual cortex in the blind follows retinotopic organization principles. Brain 138:1679–1695.

Sunness JS, Liu T, Yantis S (2004): Retinotopic mapping of the visual cortex using functional magnetic resonance imaging in a patient with central scotomas from atrophic macular degeneration. Ophthalmology 111:1595–1598.

Timberlake GT, Peli E, Essock EA, Augliere RA (1987): Reading with a macular scotoma. II. Retinal locus for scanning text. Invest Ophthalmol Vis Sci 28:1268– 1274.

Tootell RB, Reppas JB, Dale AM, Look RB, Sereno MI, Malach R, Brady TJ, Rosen BR (1995): Visual motion aftereffect in human cortical area MT revealed by functional magnetic resonance imaging. Nature 375:139–141.

Toyoizumi T, Kaneko M, Stryker MP, Miller KD (2014): Modeling the dynamic interaction of Hebbian and homeostatic plasticity. Neuron 84:497–510.

Tropea D, Kreiman G, Lyckman A, Mukherjee S, Yu H, Horng S, Sur M (2006): Gene expression changes and molecular pathways mediating activity-dependent plasticity in visual cortex. Nat Neurosci 9:660–668.

Turrigiano GG (1999): Homeostatic plasticity in neuronal networks: the more things change, the more they stay the same. Trends Neurosci 22:221–227.

Wolbers T, Wiener JM, Mallot HA, Büchel C (2007): Differential recruitment of the hippocampus, medial prefrontal cortex, and the human motion complex during path integration in humans. J Neurosci 27:9408–9416.

Yan W, Peng Y-R, van Zyl T, Regev A, Shekhar K, Juric D, Sanes JR (2020): Cell atlas of the human fovea and peripheral retina. Sci Rep 10:9802.

Yu H-H, Verma R, Yang Y, Tibballs HA, Lui LL, Reser DH, Rosa MGP (2010): Spatial and temporal frequency tuning in striate cortex: functional uniformity and specializations related to receptive field eccentricity. Eur J Neurosci 31:1043–1062.

Zarbin MA (2004): Current concepts in the pathogenesis of age-related macular degeneration. Arch Ophthalmol 122:598–614.

Zhang LI, Bao S, Merzenich MM (2001): Persistent and specific influences of early acoustic environments on primary auditory cortex. Nat Neurosci 4:1123–1130.

Zhen Z, Kong X-Z, Huang L, Yang Z, Wang X, Hao X, Huang T, Song Y, Liu J (2017): Quantifying the variability of scene-selective regions: Interindividual, interhemispheric, and sex differences. Hum Brain Mapp 38:2260–2275.

