## SupplementalFile for "Impact of deprivation and preferential usage on functional connectivity between early visual cortex and category selective visual regions"

**Supplementary Material**

***I. MRI DATA PROCESSING***

As described in the main text, the MRI data pre-processing was performed with FMRIPrep version 1.2.5 (https://fmriprep.org/) under the default settings [^57^](https://sciwheel.com/work/citation?ids=6124019&pre=&suf=&sa=0). The information below is a detailed description of the methods taken from the automatically generated output of the fMRIPrep pipeline:

*Anatomical data preprocessing*

T1-weighted images were corrected for intensity non-uniformity and skull-stripped. Brain surface reconstruction was performed using recon-all from FreeSurfer v6.0.0[^58^](https://sciwheel.com/work/citation?ids=893393&pre=&suf=&sa=0). Spatial normalization to the ICBM 152 Nonlinear Asymmetrical template version 2009c was performed through nonlinear registration with antsRegistration (ANTs 2.2.0[^59^](https://sciwheel.com/work/citation?ids=1353025&pre=&suf=&sa=0)), using brain-extracted versions of both T1w volume and template. Brain tissue segmentation of cerebrospinal fluid (CSF), white-matter (WM) and gray-matter (GM) was performed on the brain-extracted T1w using FSL 5.0.9[^60^](https://sciwheel.com/work/citation?ids=1071419&pre=&suf=&sa=0).

*Functional data preprocessing*

Functional MRI data preprocessing in fMRIPrep was carried out as follows: First, a reference volume and its skull-stripped version were generated using fMRIPrep. The BOLD reference was then co-registered to the T1w reference using bbregister (FreeSurfer) which implements boundary-based registration[^61^](https://sciwheel.com/work/citation?ids=559299&pre=&suf=&sa=0). Co-registration was configured with nine degrees of freedom to account for distortions remaining in the BOLD reference. Head-motion parameters with respect to the BOLD reference (transformation matrices, and six corresponding rotation and translation parameters) are estimated before any spatiotemporal filtering[^62^](https://sciwheel.com/work/citation?ids=8414456&pre=&suf=&sa=0). BOLD runs were slice-time corrected using 3dTshift from AFNI 20160207 [^63^](https://sciwheel.com/work/citation?ids=1524112&pre=&suf=&sa=0). The BOLD time-series (including slice-timing correction) were resampled onto their original, native space by applying a single, composite transform to correct for head-motion and susceptibility distortions. These resampled BOLD time-series will be referred to as preprocessed BOLD in original space, or just preprocessed BOLD. The BOLD time-series were resampled to MNI152NLin2009cAsym standard space, generating a preprocessed BOLD run in MNI152NLin2009cAsym space. First, a reference volume and its skull-stripped version were generated using fMRIPrep. Several confounding time-series were calculated based on the preprocessed BOLD: framewise displacement (FD), DVARS, and three region-wise global signals. FD and DVARS are calculated for each functional run, both using their implementations in *Nipype* 1.1.6[^64^](https://sciwheel.com/work/citation?ids=2539698&pre=&suf=&sa=0) following the definitions by[^65^](https://sciwheel.com/work/citation?ids=24074&pre=&suf=&sa=0). The three global signals are extracted within the CSF, the WM, and the whole-brain masks. Additionally, a set of physiological regressors were extracted to allow for component-based noise correction[^66^](https://sciwheel.com/work/citation?ids=2716556&pre=&suf=&sa=0). Principal components are estimated after high-pass filtering the preprocessed BOLD time-series (using a discrete cosine filter with 128s cut-off) for the two CompCor variants: temporal (tCompCor) and anatomical (aCompCor). Six tCompCor components are then calculated from the top 5% variable voxels within a mask covering the subcortical regions. This subcortical mask is obtained by heavily eroding the brain mask, which ensures it does not include cortical GM regions. For aCompCor, six components are calculated within the intersection of the aforementioned mask and the union of CSF and WM masks calculated in T1w space, after their projection to the native space of each functional run (using the inverse BOLD-to-T1w transformation). The head-motion estimates calculated in the correction step were also placed within the corresponding confounds file. All resamplings can be performed with a single interpolation step by composing all the pertinent transformations (i.e. head-motion transformation matrices, susceptibility distortion correction when available, and co-registrations to anatomical and template spaces). Gridded (volumetric) resamplings were performed using antsApplyTransforms (ANTs), configured with Lanczos interpolation to minimize the smoothing effects of other kernels[^67^](https://sciwheel.com/work/citation?ids=6017381&pre=&suf=&sa=0). Non-gridded (surface) resamplings were performed using mri_vol2surf (FreeSurfer). For further details of the pipeline, including the software packages utilized by FMRIPrep for each preprocessing step, please refer to the online documentation: [https://fmriprep.readthedocs.io/en/](https://fmriprep.readthedocs.io/en/1.0.3/)1.2.5/.

*Motion artifact scrubbing*

Motion artifacts during fMRI scanning are known to produce substantial effects on functional connectivity data [^68,69^](https://sciwheel.com/work/citation?ids=1374987,1475085&pre=&pre=&suf=&suf=&sa=0,0). In order to mitigate such confounds, we performed motion scrubbing on the preprocessed data from fMRIPrep using the XCP engine workflow [^70^](https://sciwheel.com/work/citation?ids=6033560&pre=&suf=&sa=0) using a framewise displacement threshold of 0.5mm. For more information on the scrubbing procedure, see online documentation: https://xcpengine.readthedocs.io/modules/regress.html#temporal-censoring. Following preprocessing, the data were converted into a template space (fsLR_32k cifti-space) using a combination of *Ciftify* [^71^](https://sciwheel.com/work/citation?ids=7441300&pre=&suf=&sa=0) and the *Connectome Workbench* [^72^](https://sciwheel.com/work/citation?ids=1352575&pre=&suf=&sa=0).

**II. Supplementary Figures**

The figures below show the individual microperimetry images used to generate the individualized surface-based, cortical regions of interest (also shown). As summarized in the methods section: microperimetry was first on each eye separately (top two panels). Images from the left and right eyes were subsequently combined in a single image (center panel). Using this images of the full visual field, regions of interest were localized in visual cortex (bottom) for each subject.

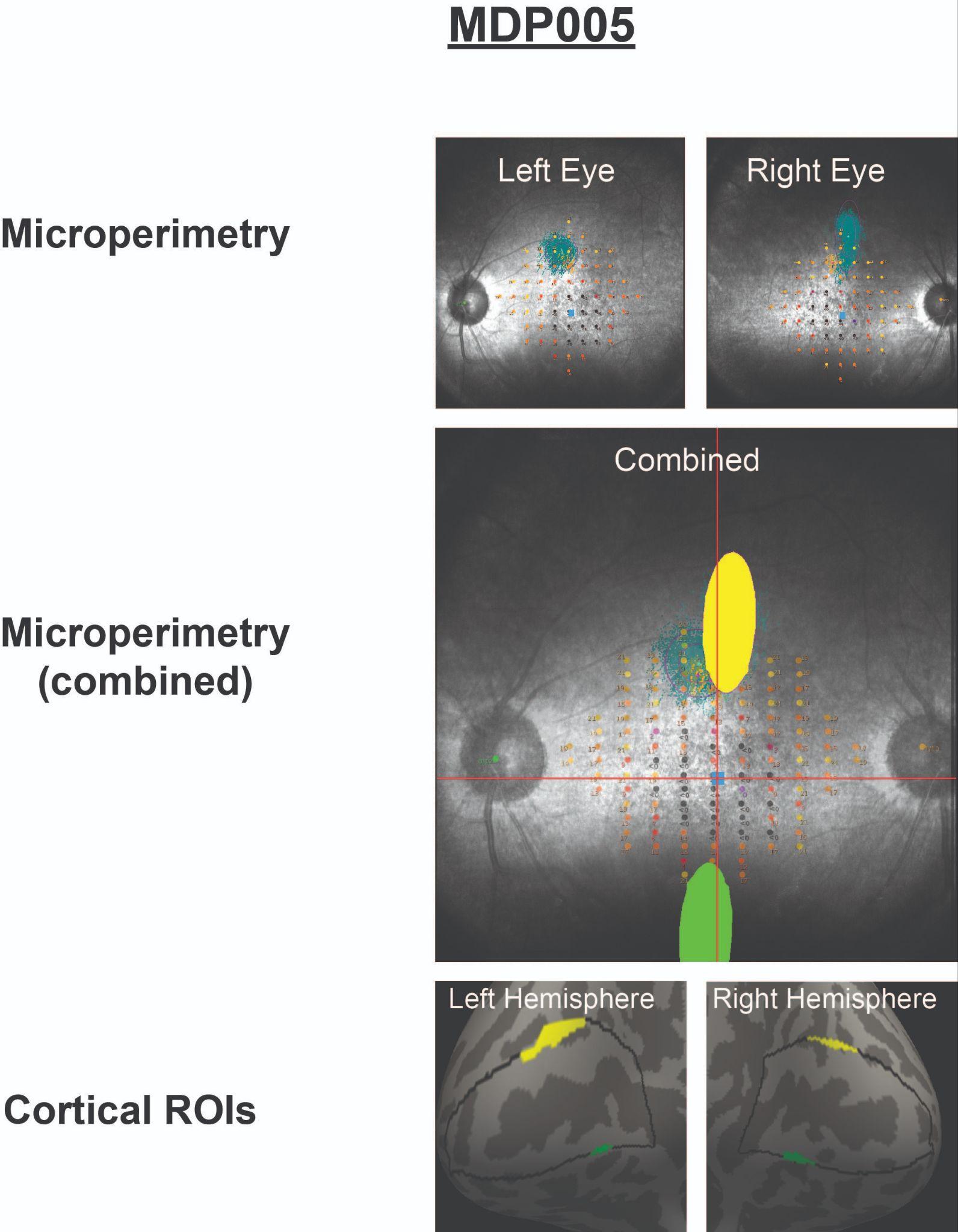

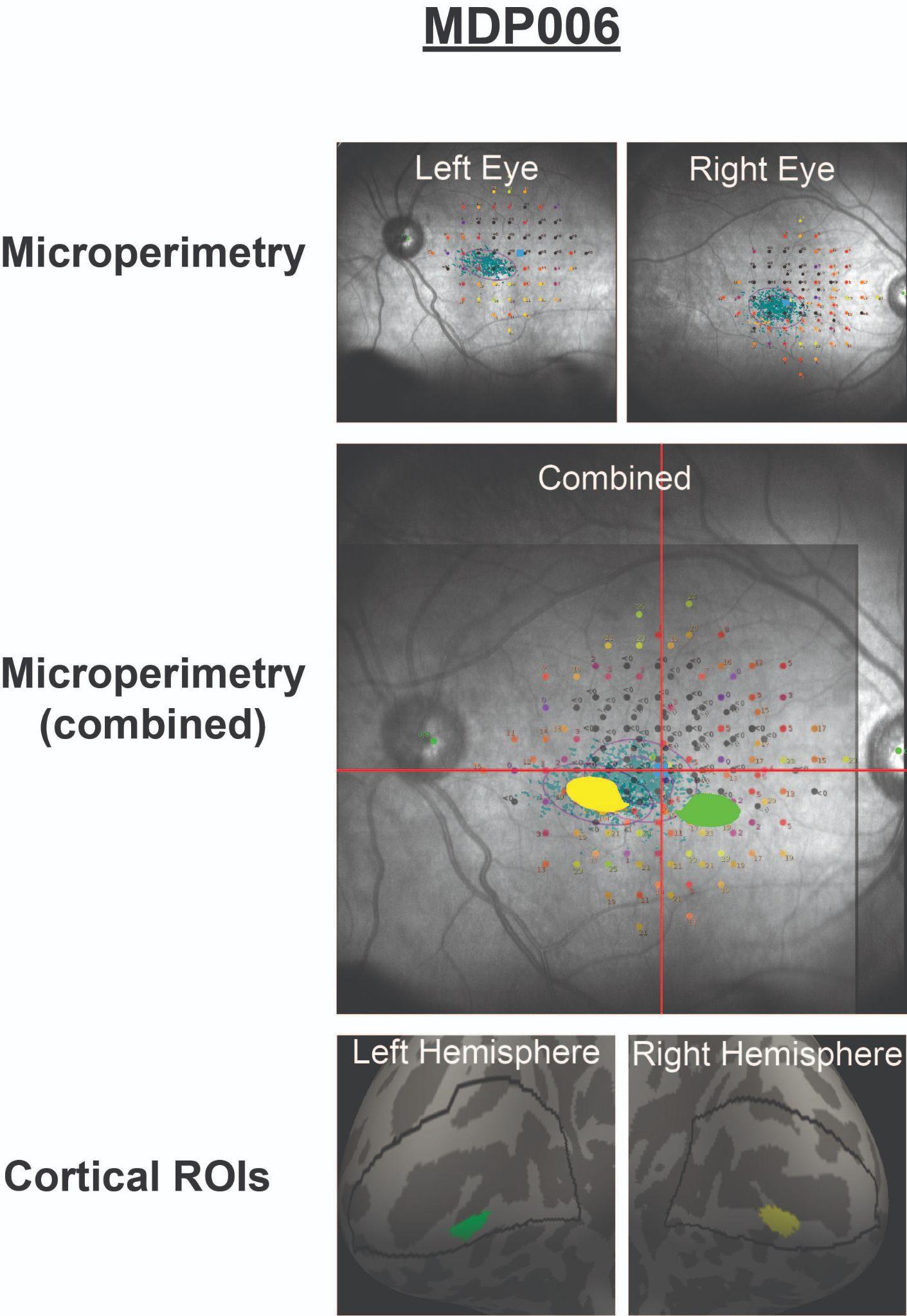

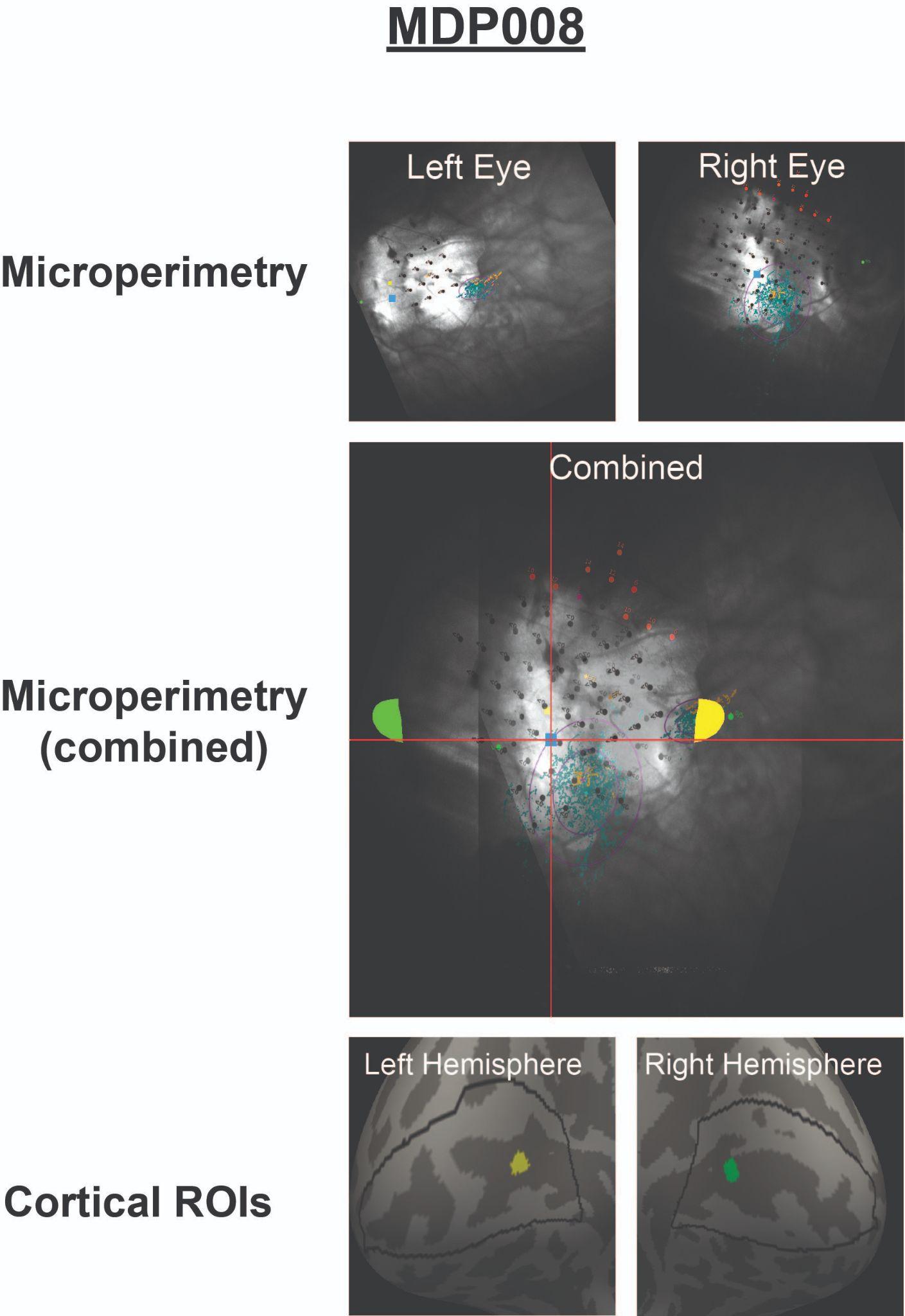

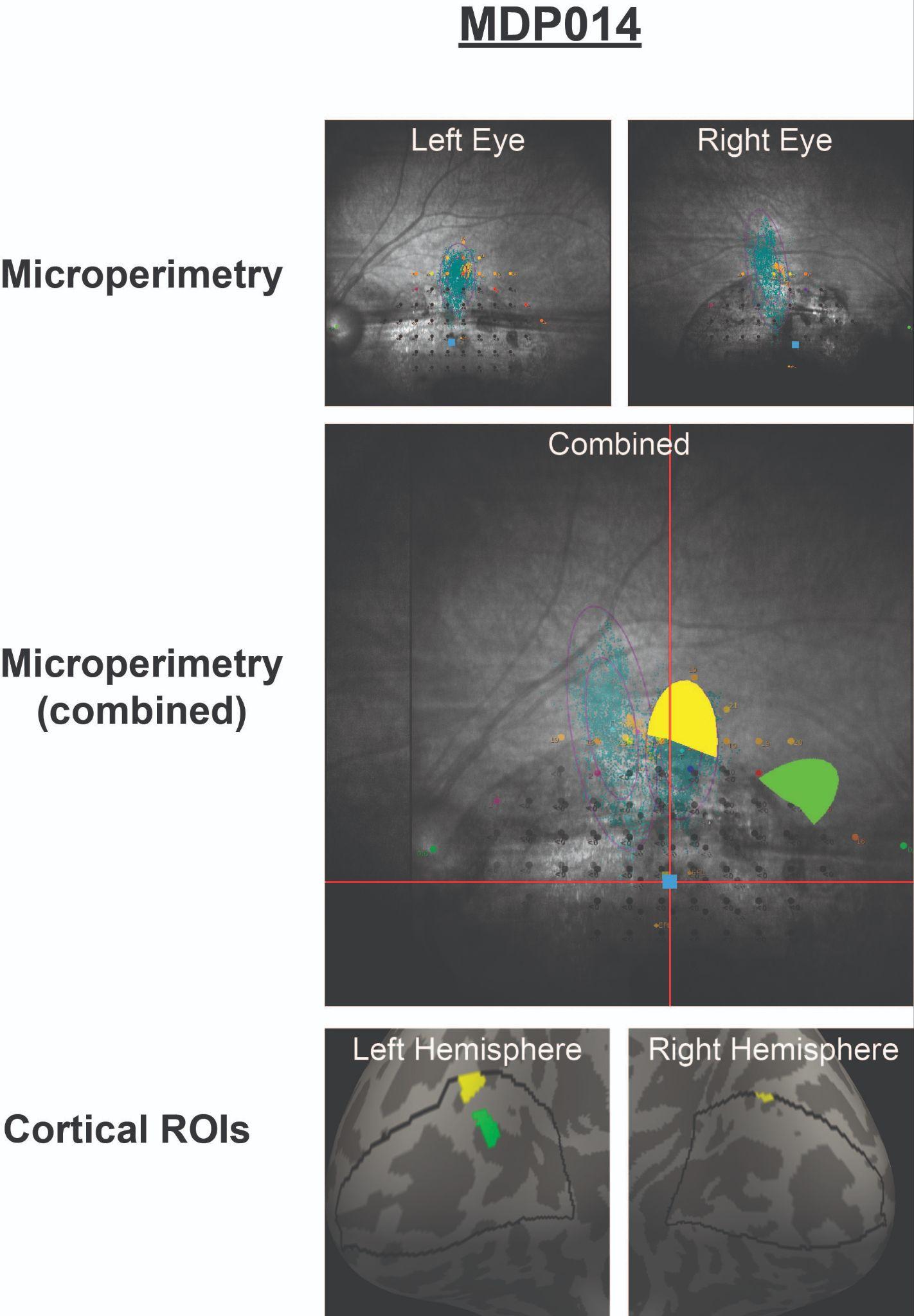

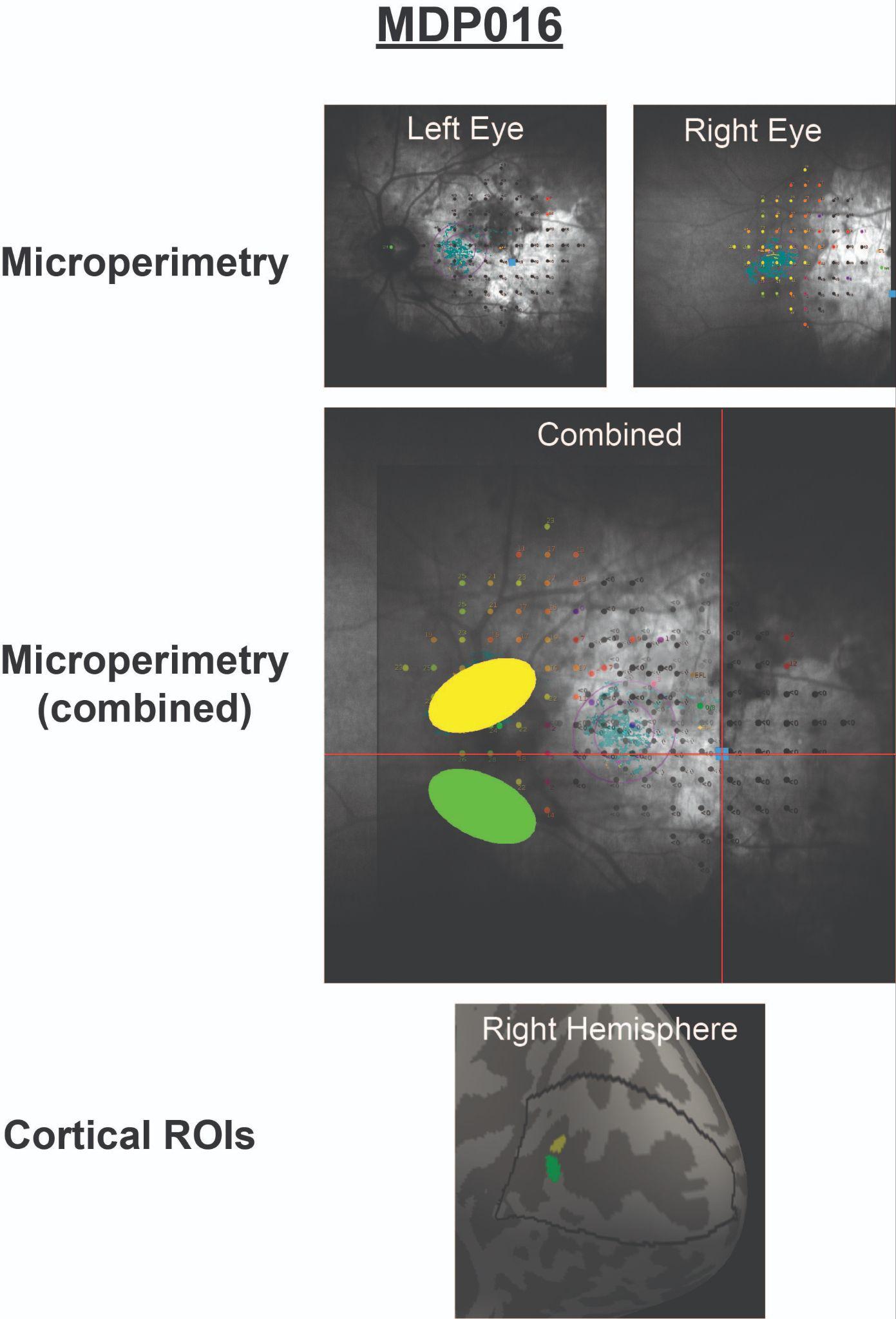

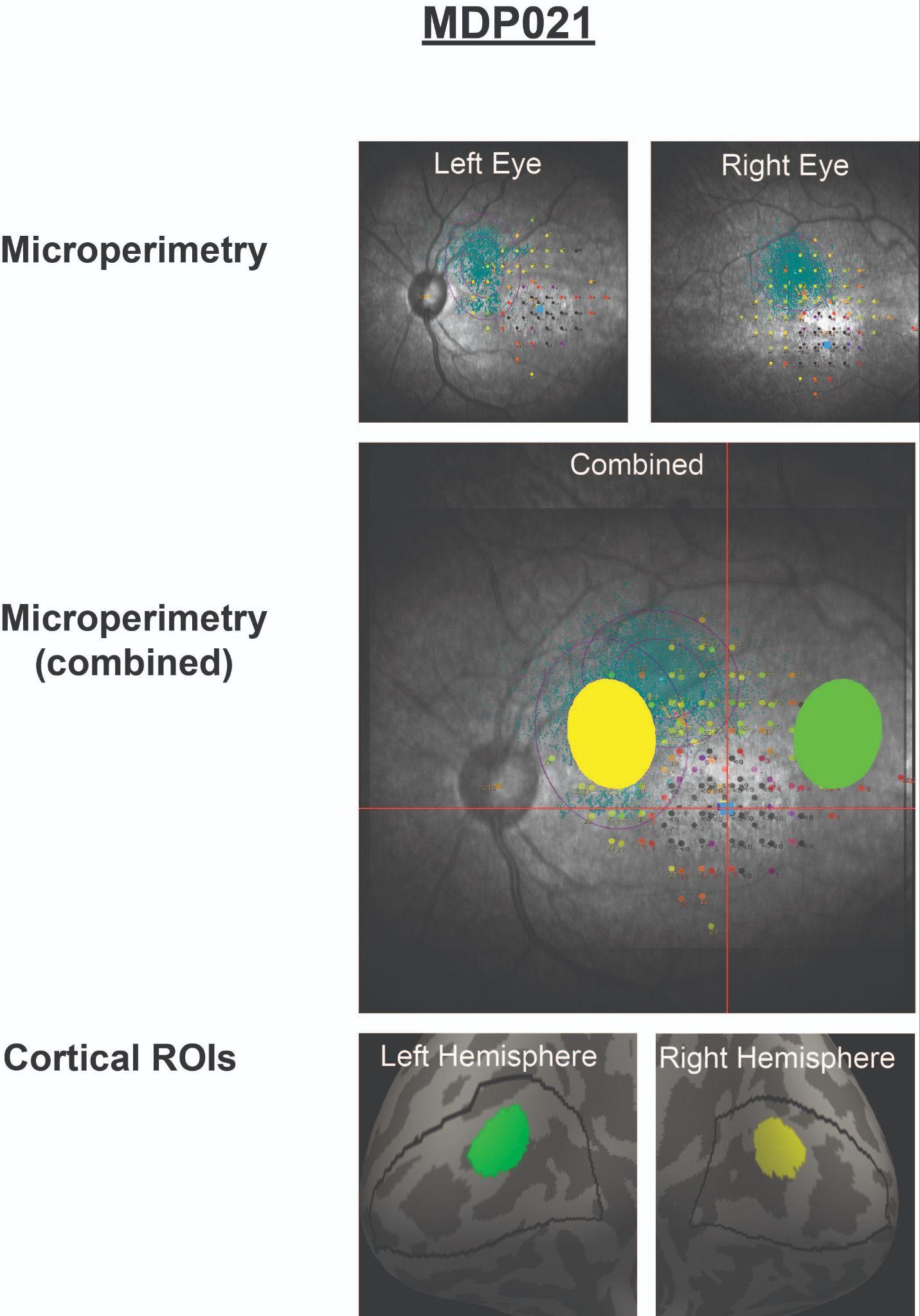

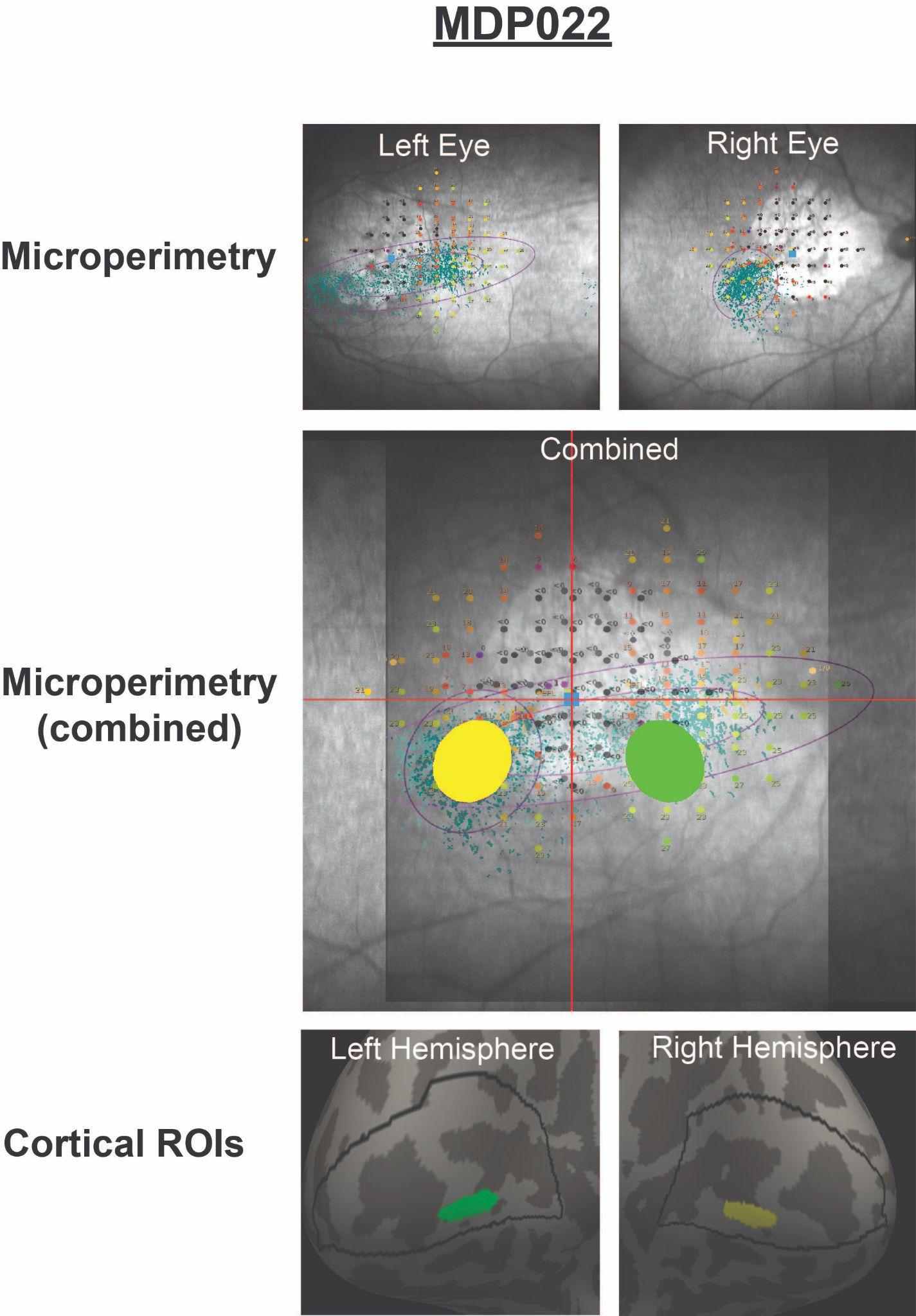

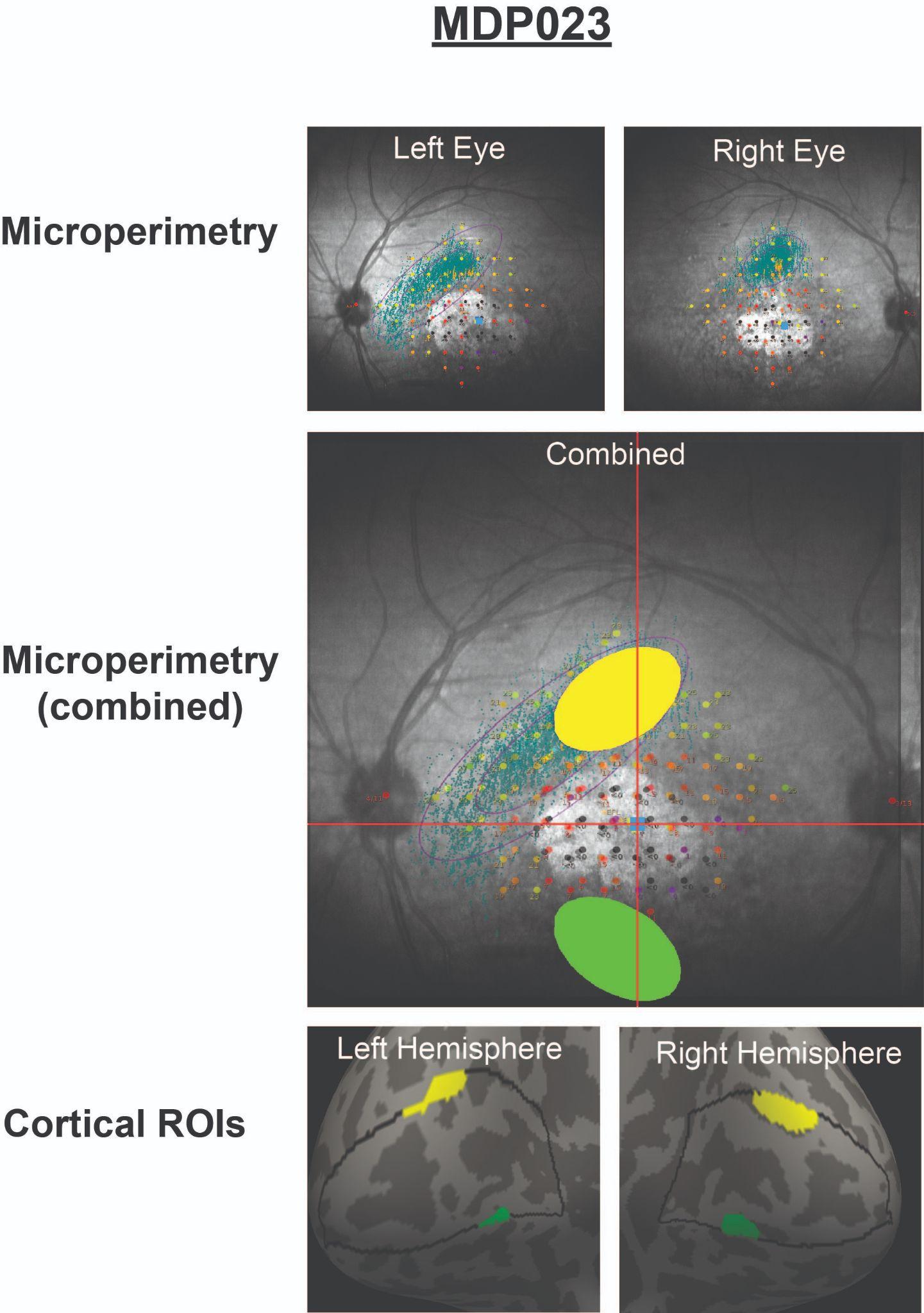

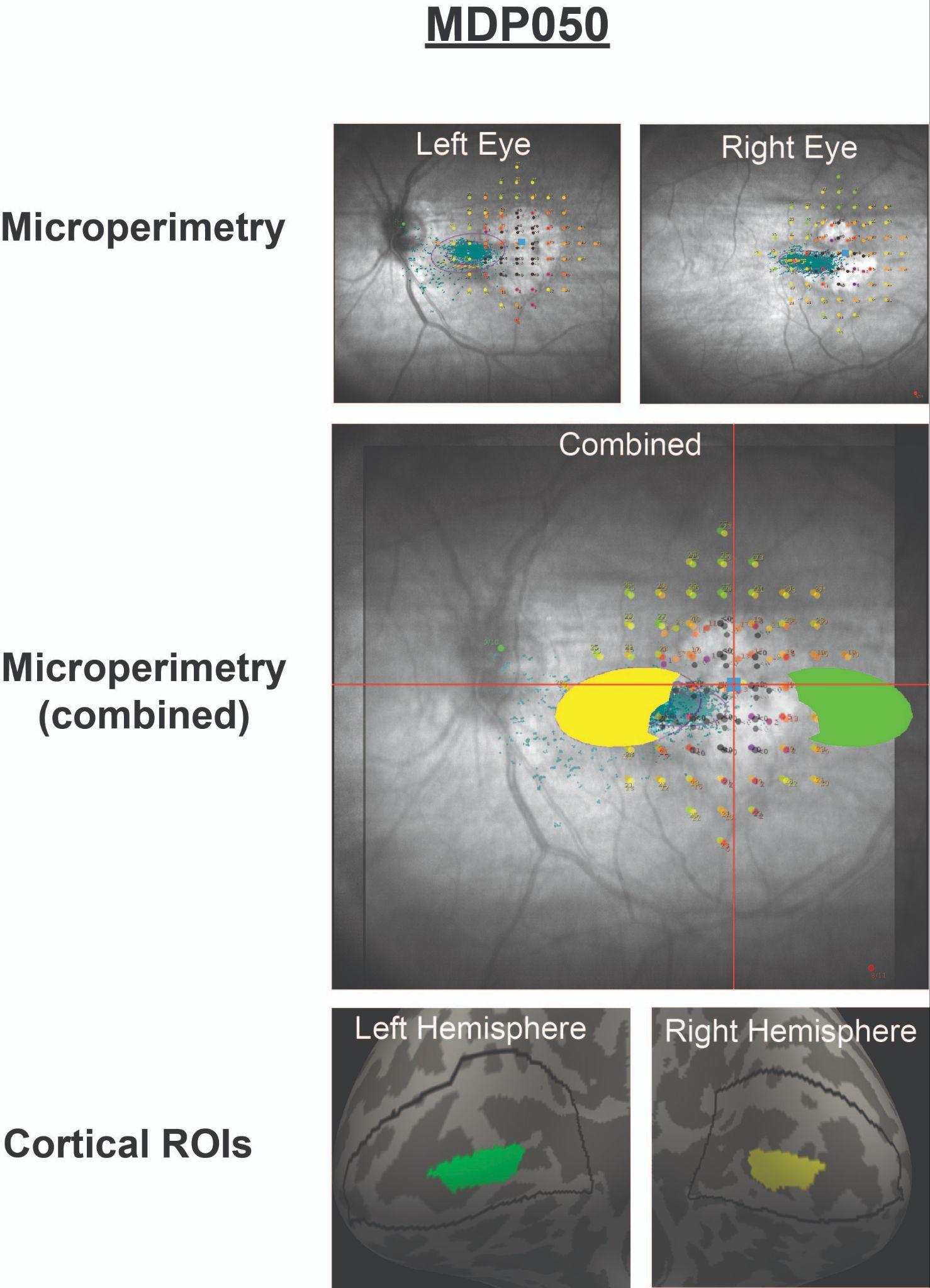

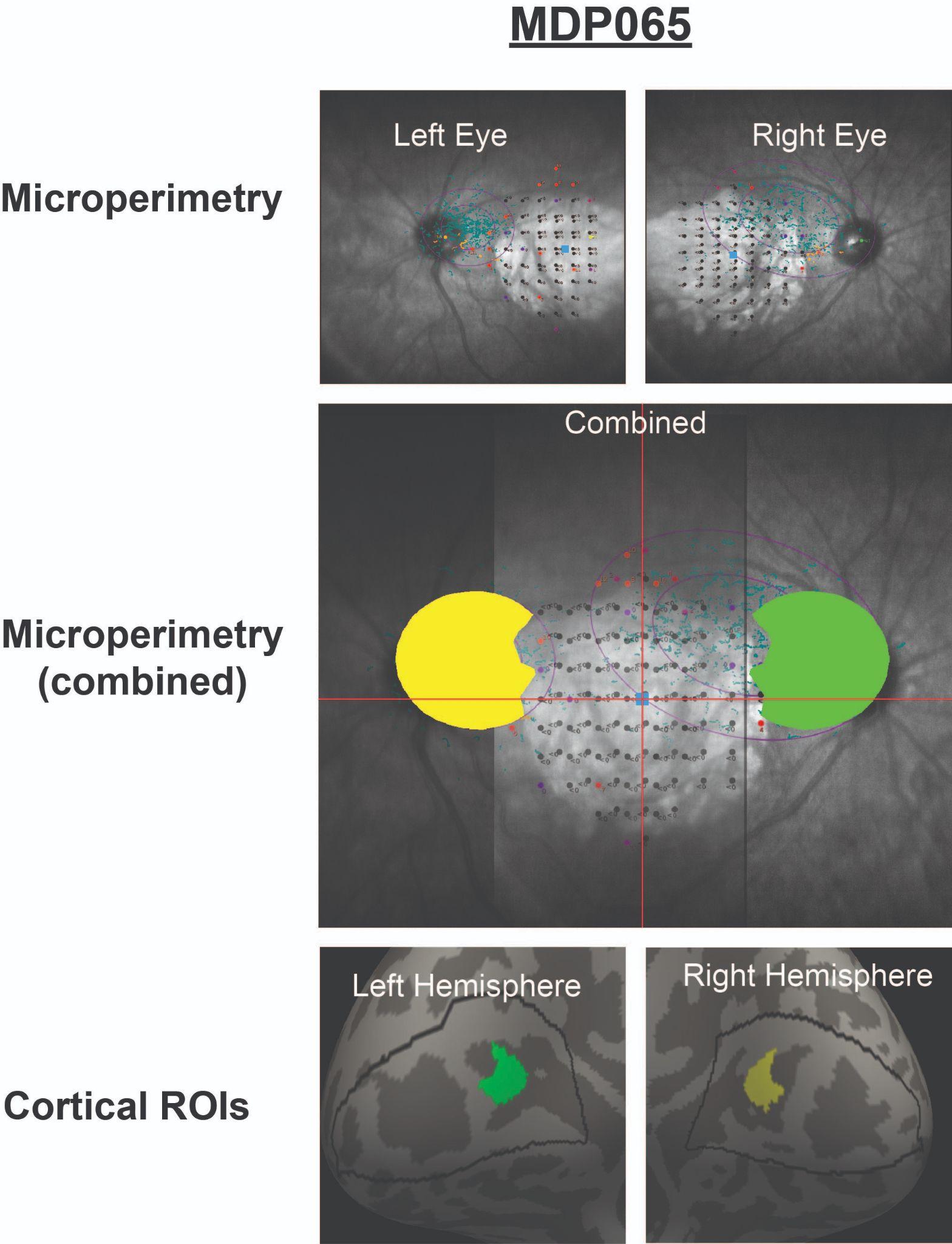

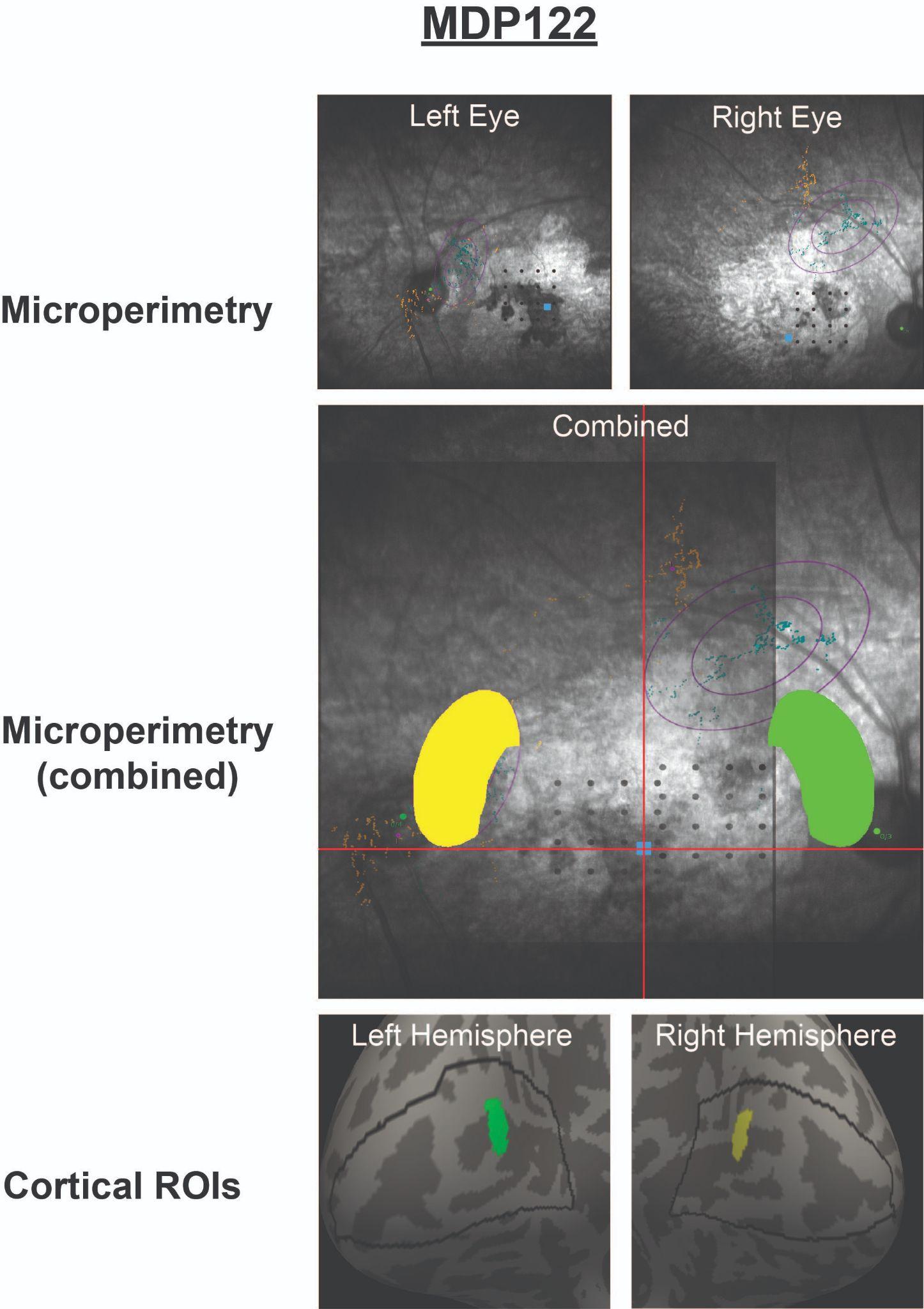
